# TDP-43 self-assembly is regulated by its disordered NLS-region

**DOI:** 10.64898/2026.06.06.730435

**Authors:** Saskia Hutten, Emre Pekbilir, Benjamin Bourgeois, Xiaofei Ping, Christoph Rickert, Nele Kuhr, Nora Knabe, Simone Mosna, Tobias Madl, Lukas S. Stelzl, Dorothee Dormann

## Abstract

Cytosolic inclusions of the RNA-binding protein TDP-43 are a pathological hallmark of several neurodegenerative diseases, such as amyotrophic lateral sclerosis and frontotemporal dementia. Cellular or animal model systems often use TDP-43 mutated in its nuclear localization signal (NLS) to study its cytosolic mislocalization and aggregation. Here we show that the disordered NLS-region, in particular the basic amino acids, are crucial for self-assembly of full-length TDP-43 across size scales, ranging from small clusters to visible condensates and aggregates. Molecular dynamics simulations and NMR studies suggest that the NLS-region engages in inter-chain interactions with C-terminal aromatic residues as well as RRM1 and NTD interactions. We further demonstrate that a minimal NLS mutation (K82A) preserves TDP-43 condensation in vitro and in cells, while commonly used NLS mutations yield partially or strongly reduced self-assembly behaviors. Our data highlight TDP-43 K82A as ideal model system to study cytosolic TDP-43 aggregation in cell and animal models.

## Introduction

The principal component of the cytosolic aggregates in brains of patients suffering from amyotrophic lateral sclerosis (ALS), frontotemporal dementia (FTD) or limbic-predominant age-related TDP-43 encephalopathy (LATE) is TAR DNA-binding protein of 43 kDa (TDP-43)^1,2^. TDP-43 is a ubiquitously expressed RNA-binding protein (RBP) and has a central role in RNA metabolism^3–5^. It consists of several folded domains: an N-terminal domain (NTD) involved in physiological head-to-tail dimerization^6^, and two central RNA-binding motifs (RRM) that mediate binding to GU-rich RNA^7,8^. At its C-terminus, TDP-43 has a largely disordered low-complexity domain (LCD), which harbors most of the known ALS-associated point mutations^9^.

TDP-43 continuously shuttles between the nucleus and cytoplasm^10,11^. However under physiological conditions, it localizes predominantly to the nucleus due to the presence of a bipartite classical nuclear localization sequence (cNLS)^12,13^. This cNLS is situated in the linker region between the NTD and RRM1 and consists of two clusters of basic amino acids, of which in particular the first cluster mediates binding to its cognate import receptor Importin α/β (Impα/β)^14^. Cellular stress can cause TDP-43 to become mislocalized to the cytoplasm, where it associates with stress-induced biomolecular condensates, so called stress granules (SGs)^15–20^. SGs might be directly linked to TDP-43 aggregation, as RBP inclusions in ALS and FTD patients frequently contain proteins found in SGs^15,18,21,22^. Whether SGs are indeed precursors of pathological TDP-43 aggregates is still under debate. While some report formation of TDP-43 aggregates independently of SGs^23–27^, others have suggested that SGs act as crucibles of TDP-43 aggregation in the cytoplasm^15,18,28–30^.

SGs are coacervates of protein and RNA that assemble through phase separation, a process by which individual macromolecules above a critical concentration de-mix from a homogeneous solution into microscopically visible condensates^31–34^. Like other proteins found in pathological inclusions or SGs, TDP-43 has been shown to phase separate and self-assembly into homotypic condensates *in vitro*^35,36^, yet the underlying inter- and intramolecular interactions are still not completely understood. The largely disordered C-terminal LCD can condense on its own^37^ and the involved residues and features have been extensively studied. In several publications, the Fawzi lab has highlighted an important role of an α-helix in the conserved region (CR; aa 321-340)^37,38^. Additionally, aromatic and aliphatic residues in the LCD were shown to play an important role in TDP-43 LCD multimerization and phase separation^39–42^. The α-helical region and aromatic/aliphatic residues were also shown to be important for TDP-43 condensate formation in cells^43–45^. An additional pre-requisite for condensation of the full-length protein is NTD-mediated dimerization, as mutations in the NTD dimerization interface (E17/E21/Q34/R52/R55A) or phosphorylation at Ser48 abolish TDP-43 condensation both *in vitro* and in cells^6,36,46^. Recent molecular dynamics simulations of TDP-43 full-length condensates also highlight both homotypic (e.g. CR-CR) as well as heterotypic (e.g. LCD with folded domains) interactions^29,47,48^, indicating that multiple regions of TDP-43 are likely involved in TDP-43’s condensation behavior. Inter-domain interactions can also drive formation of much smaller, nano- and mesoscale oligomers or clusters of heterogenous sizes below the diffraction limit^49–51^. Whether mutations that interfere with microscopic phase separation also affect TDP-43 self-assembly at sub-critical concentrations is currently not known.

Condensation has been intimately linked to the formation of disease-linked aggregates, since numerous studies have demonstrated that liquid-like RBP droplets can transition into solid-like fibrous or amorphous RBP aggregates^52–54^. TDP-43 aggregates in ALS/FTD patients consist of a detergent-insoluble and protease-resistant amyloid-like core comprising the CR and flanking residues of the LCD^55–57^. However, it is currently still unknown how exactly TDP-43 aggregates form in disease and which sequence features drive the aggregation process.

Cellular or animal model systems to study TDP-43 aggregation often utilize point mutations in the NLS to cause cytosolic mislocalization and investigate the aggregation process in the absence of nucleocytoplasmic shuttling. Yet, whether the linker region comprising the NLS alters TDP-43 self-association is still unknown. Using *in vitro* assays with recombinant TDP-43, we find that this disordered linker region and especially the basic clusters of the cNLS are required for visible TDP-43 phase separation and cluster formation at subcritical concentrations. Molecular dynamics simulations suggest that these basic residues are involved in heterotypic domain interactions of the NLS with aromatic residues in the LCD, and NMR studies support their involvement in RRM1 and NTD interactions. A systematic comparison of different commonly used import-deficient NLS mutants demonstrates that the number of mutations in basic residues correlates with the self-assembly defect of TDP-43 *in vitro*. In cells, mutation of both basic amino acid clusters in the cNLS abolishes TDP-43 condensation and aggregation in the cytoplasm, while a less invasive mutation of just the first basic amino acid cluster causes a much more moderate alteration to the TDP-43 self-assembly behavior. In summary, our data show an important role of the NLS-region for TDP-43’s self-assembly behavior and suggest that minimal mutation of its NLS sequence at K82 is best suited for cellular and animal models aimed at studying cytoplasmic TDP-43 condensation and aggregation.

## Material and Methods

### Antibodies

The following primary and secondary antibodies were used: rabbit anti TDP-43 (proteintech 80002-1-RR or 10782-2-AP), monoclonal rat anti TDP-43 6D6 (provided by Monoclonal Antibody Core Facility, Helmholtz Munich), pSer409/410 TDP-43 (proteintech, 80007-1-RR), mouse anti GFP (proteintech # 66002-1-Ig), rabbit anti G3BP (proteintech; 13057-2-AP), mouse anti MBP (proteintech, 66003-1-Ig), mouse anti α-tubulin (proteintech, 66031-1-Ig), mouse anti GAPDH (proteintech, 60004-1-Ig); donkey anti mouse AF488 (Thermo, A-21202), donkey anti rabbit AF-Rb555 (Thermo, A-31572), donkey anti rabbit AF647 (Thermo, A-31573); donkey anti mouse IRDye 800CW (Li-cor, 926-32212); donkey anti rabbit IRDye 800CW (Licor, 926-32213), donkey anti mouse IRDye 680RD (Li-cor, 926-68072), donkey anti rabbit IRDye 680RD (Li-cor, 926-68073), goat anti rat IRDye 680RD (Li-cor, 926-68076). The monoclonal TDP-43 clone 6D6 was generated as described before^58^ and purified from cell culture supernatant using Protein A Sepharose.

### cDNA expression constructs

For recombinant TDP-43, corresponding mutations were introduced into the pJ4M-TDP-43-tev-MBP-His_6_ plasmid^36^ (Addgene #104480) either by site-directed mutagenesis or via synthetic double-stranded DNA-fragments (gBlocks gene fragments, IDT). For site directed mutagenesis, a Q5 high fidelity DNA polymerase (NEB) was used together with mutation-specific primers. gBlocks were synthesized including the desired mutations and inserted using suitable restriction enzymes. The construct for WT TDP-43 NTF (N-terminal fragment; aa 1-266)-tev-MBP-His was described previously^59^, the NTF carrying the RK6A mutation was derived by PCR from the TDP-43 (RK6A)-tev-MBP-His_6_ construct.

For generation of stable HeLa cell lines, EGFP-tagged expression constructs for TDP-43 K82A and 83AAA (K82/R83/K84A) were generated in the pcDNA5-FRT/TO-EGFP-TDP-43 backbone^60^. For RK6A, the GFP-TDP-43 ORF was PCR-amplified from a GFP-TDP-43 RK6A^35^ construct and inserted into the pcDNA5-FRT/TO-backbone cut with HindIII and BamHI using Gibson assembly. For K82A, the mutation was introduced in pcDNA5-FRT/TO-EGFP-TDP-43 WT using site directed mutagenesis. The cDNA sequence for TDP-43 83AAA was PCR amplified from a construct, in which the mutation had been inserted using QuikChange mutagenesis (Stratagene)^15^. Note, all these constructs were rendered resistant against the siRNA used to knockdown endogenous TDP-43 (see *Maintenance, cDNA transfection, siRNA and stress treatment).* K82A and RK6A mutants of the GCR_2_-EGFP_2_-TDP-43 reporter^10^ were generated by mutagenesis carrying primer with the desired mutations. All constructs were verified by Sanger sequencing.

### Cell Culture

#### Generation of stable cell lines

Stable HeLa cells expressing the different EGFP-TDP-43 variants in a doxycycline-inducible manner were generated using the Flp-In T-Rex system. The stable cell line for EGFP-TDP-43 WT has been described previously^60^. To generate HeLa lines stably expressing EGFP-TDP-43 K82A or 83AAA, Flp-In T-Rex HeLa cells (gift by Christian Behrends, LMU Munich) were co-transfected with pcDNA5-FRT/TO-EGFP-TDP-43 K82A or 83AAA constructs, respectively, with a 7-fold excess of the pOG44 Flp-recombinase expression plasmid (Invitrogen). Subsequently, cells were selected using Hygromycin (150µg/ml) and Blasticidin (10 µg/ml). Surviving cells were FACS-sorted using a Bigfoot cell sorter (Invitrogen, IMB Mainz Flow Cytometry core facility) to obtain single cell clones. Clones with homogenous and comparable expression for EGFP-TDP-43 were chosen for final experiments. The cell line for GFP-TDP-43 RK6A was generated by co-transfection of 10-fold excess of pOG44 over pcDNA5-FRT/TO-EGFP-TDP-43 RK6A and subsequently selected using antibiotics as described above. The final cell population was FACS-sorted based on EGFP fluorescence intensity upon induction with 1 µg/ml doxycycline and individual clones screened for homogeneity of EGFP-expression by Opera high-throughput imaging. For all experiments, EGFP-TDP-43 expression was induced upon addition of 20 ng/ml doxycycline (dox) for at least 18h.

#### Maintenance, cDNA transfection, siRNA and stress treatment

All cell lines were maintained in a humified chamber with 5% CO_2_ at 37°C. HeLa cells were grown using DMEM high glucose GlutaMAX (Invitrogen), supplemented with 10% fetal bovine serum (FBS; Invitrogen) and 50 µg/ml gentamycin. For growth of stable HeLa Flp-In lines, DMEM high glucose GlutaMAX (Invitrogen) was supplemented with 10% Tet-approved FBS (PAN or Invitrogen), 50 µg/ml gentamycin, 150 µg/ml Hygromycin and 10 µg/ml Blasticidin. cDNA and siRNA transfections were carried out using Lipofectamine 2000 or Lipofectamine RNAiMaXX (Invitrogen), respectively, omitting any antibiotics from time of transfection until analysis. For DNA transfections, medium was exchanged ca. 4-6h after cDNA transfections to reduce cellular stress caused by the transfection reagent. Unless stated otherwise, knockdown of endogenous TDP-43 in stable HeLa FlpIn GFP-TDP-43 cells was achieved by transfection with 10 nM siRNA (target sequence: 5’-AATAACGAGGGTAACACTGGG-3’; Biomers) and cells analyzed 48-58h post-transfection. Oxidative stress was induced by addition of 0.5 mM Na-arsenite (Sigma) to the medium for 1h.

### Recombinant protein purification

Generally, bacterial cells were lysed with the help of lysozyme (0.1 mg/ml) and sonication. Bacterial lysates were subsequently cleared by centrifugation (20,000g 1h 4°C) before the affinity purification step. Finally, protein concentrations were calculated by A280 measurements applying the respective molecular weight and extinction coefficient (ε) predicted by the ProtParam tool for each protein. Proteins were aliquoted, snap-frozen in liquid nitrogen and stored at −80°C.

#### TDP-43-tev-MBP-His_6_

All TDP-43-tev-MBP-His_6_ variants were purified as reported previously^35,36^ with minor modifications. In general, expression was performed in *E. coli* BL21-DE3 Rosetta 2 cells using

0.5 mM IPTG overnight at 16°C. Cells were lysed in lysis buffer (50 mM Tris pH 8.0, 1 M NaCl, 10% (v/v) glycerol, 10 mM imidazole, 4 mM β-mercaptoethanol (β-ME) and 1 µg/ml each of aprotinin, leupeptin hemisulfate and pepstatin A supplemented with 0.1 mg/ml RNase A. TDP-43-tev-MBP-His_6_ was purified from the lysate by Ni-NTA beads (Qiagen or Thermo) and eluted using 300 mM imidazole in lysis buffer. Finally, pure protein was obtained by size exclusion chromatography (SEC; HiLoad 16/600 Superdex 200 pg, GE Healthcare) in SEC-buffer (50 mM Tris pH 8.0, 300 mM NaCl, 5% glycerol (v/v) supplemented with 2 mM TCEP). Purified TDP-43 variants were concentrated using Amicon ultra centrifugal filters (30K MWCO). The A260/280 ratio was reliably found to be ≤0.6.

#### TDP-43 NTF (aa 1-266)-tev-MBP-His_6_

For expression of recombinant uniformly ^15^N-labeled WT and RK6A TDP43 NTF (aa 1-266) - tev-MBP-His_6_, the respective bacterial expression constructs were transformed into *E. coli* BL21-DE3 Star. 10 ml of liquid preculture were then transferred in minimal medium supplemented with 6 g of unlabeled glucose (Roth) and 1 g of ^15^NH_4_Cl (Eurisotop). Cells were grown to an optical density (OD600) of 0.8 and protein expression was induced by addition of 0.5 mM IPTG followed by incubation at 20°C for 16 hours. Cell pellets were harvested and sonicated in resuspension buffer (50 mM Tris-HCl pH 7.5, 150 mM NaCl, 20 mM imidazole, 2 mM TCEP). Recombinant proteins were then purified using Ni-NTA agarose (Qiagen) and the MBP-His_6_ tag was cleaved off by adding 2 (w/w) % His_6_-tagged TEV protease for 16 h at 4 °C. Untagged TDP43 WT and mutant NTFs were subsequently isolated by a second Ni-NTA affinity purification. A final size exclusion chromatography purification step was performed in the buffer of interest on a gel filtration column (Superdex 75; Cytiva).

#### His_6_-Tev protease

Expression of His_6_-Tev protease was induced in E. coli BL21-DE3 Star by 1 mM IPTG o/n at 20°C. Cells were lysed in lysis buffer (50 mM Tris pH8, 200 mM NaCl, 10% glycerol (v/v), 20 mM imidazole, 4 mM β-ME) supplemented with 0.1 mg/ml RNase A. His-Tev was affinity purified using Ni-NTA agarose (Qiagen or Thermo), washed with lysis buffer and once with lysis buffer containing 1 M NaCl. Elution was achieved by 400 mM imidazole in lysis buffer and the protein subsequently dialyzed overnight into storage buffer (20 mM Hepes pH 7.4, 150 mM NaCl, 20% glycerol, 2 mM DTT).

#### His_6_-S-Impβ, His_6_-Impα3 and RanQ69L

Expression and purification of His*_6_*-S-Impβ^61^, His*_6_*-Impα3^61^ and RanQ69L^62^ was performed as described previously.

### Phase separation Assays

Prior to all phase separation assays, recombinant TDP-43 was centrifuged after thawing to remove any preformed aggregates and its concentration was verified using A280 measurements as described above (*Recombinant protein purification*).

#### Protein sequence analysis

Net-Charge-Per-Residue (NCPR) was calculated using the online CIDER tool^63^. Order-Disorder predictions were performed using IUPRED3 and PrDOS^64,65^.

#### Sedimentation assay (solubility assay)

For analysis of solubility of recombinant TDP-43, 5 µM TDP-43-tev-MBP-His_6_ was cleaved for 1h at room temperature (RT) by addition of His_6_-Tev protease in condensation buffer (20 mM Hepes pH 7.5, 150 mM NaCl (final), 2 mM DTT) followed by centrifugation at 21,000g for 15 min at 4°C. Equal volumes of supernatant (S) and pellet (P) were analyzed by western blot analysis using TDP-43 specific antibodies as indicated in the respective figure legends.

#### Turbidity assay

Turbidity measurements were performed in 20 µl reaction volume in technical duplicates in 384- well low binding plates (Greiner). 10 µM TDP-43-tev-MBP-His_6_ was cleaved using 3.2 µM His_6_-Tev protease in condensation buffer and turbidity measured at 350 nm in a SpectraMax ID5 plate reader.

#### Microscopic inspection of condensates

For visual inspection of condensate morphology using phase contrast microscopy, all reactions were performed using Ibidi µ-Slide 18-well flat chambers (ibidi, #81826). TDP-43-tev-MBP-His_6_ at indicated concentrations was incubated with His_6_-Tev protease at a fixed ratio (2.5x molar excess TDP-43) for ca. 1h before image acquisition using a Zeiss Observer. For analysis of the impact of RNA on TDP-43 condensate formation, increasing amounts of a Cy5-labelled RNA corresponding to the autoregulatory element (ARE) in the TDP-43 mRNA (5’-Cy5-GAGAGAGCGCGUGCAGAGACUUGGUGGUGCAUAA-3’; IDT) were added to the reaction before adding His_6-_Tev protease. Images of fluorescent RNA were acquired using structural illumination (Zeiss Apotome; see below).

#### Solubility upon sarkosyl extraction

To generate aggregated forms of recombinant TDP-43, we utilized a published protocol^35,66^ with slight modifications. In brief, 5 µM recombinant TDP-43-tev-MBP-His_6_ variants (WT, K82A, 83AAA, RK6A) in aggregation buffer (50 mM Tris pH 8.0, 250 mM NaCl, 5% glycerol, 5% sucrose, 150 mM imidazole, 2 mM DTT) were cleaved by addition of 1.5 µM His_6-_Tev- protease in protein low binding tubes (Sarstedt). After incubation for 30 minutes at room temperature, samples were agitated by shaking at 1000 rpm at room temperature (22°C) for 30 minutes. Samples were then either directly processed (D0) or incubated for three days at RT without agitation (D3). To address detergent solubility, samples were incubated at D0 or D3, respectively, with 1 % (final) Sarkosyl (N-Lauroylsarcosine sodium salt, Sigma) with slight agitation (300 rpm) at 37 °C for 1 hour, followed by centrifugation at 21,000g for 15 minutes at 4°C. Equal amounts of supernatant (S) and pellet (P) fraction were subsequently analyzed by western blot using a TDP-43-specific antibody (proteintech # 80002-1-RR).

#### Dynamic Light Scattering (DLS)

To detect small assemblies of recombinant TDP-43 below the visible range, we performed DLS measurements of TDP-43 at subcritical concentrations (0.7 µM) in condensation buffer 1h after Tev protease cleavage using a Zetasizer Ultra (Malvern Pananalytical). Importantly, all buffers were filtered using 0.22 µm membranes (Millex-GV PVDF or Millex-GP PES, Milipore). Measurements were conducted in 100 µl volume using low volume disposable cuvettes (Brandt) with the following settings (T: 20°C, equilibration time: 30 sec; angle: 173°, dispersant: H_2_O; material: protein) utilizing auto mode of ZS Xplorer Software (Malvern Pananalytical) and the in-built “General Purpose” model for data analysis.

### Electrophoretic Mobility Assay (EMSA)

For analysis of RNA-binding capacity of recombinant TDP-43 WT and mutants, 2 nM of IR700-labelled ARE-RNA probe (5’-IR700-GAGAGAGCGCGUGCAGAGACUUGGUGGUGCAUA A-3’; metabion) were diluted in binding buffer (20 mM HEPES pH 7.5, 150 mM NaCl, 5 mM MgCl_2_, 0.01 µg/µl yeast tRNA, 0.1 µg/µl BSA, 2 mM DTT, 1 U/µl RNase inhibitor (Thermo Fisher)) and mixed with varying amounts of TDP-43-tev-MBP-His_6_ WT, 4FL, delNLS, and RK6A. Samples (10 µl) were incubated for 30-45 min at RT to allow for RNA-binding and after addition of glycerol (6% (v/v) final; corresponding to 1/6^th^ of final sample volume) analyzed by 6% non-denaturing polyacrylamide gel electrophoresis. Gels were run in 0.5x TBE at 100 V for 40 min at RT before imaging with a Licor Odyssey M imaging system.

### Pulldown (PD) analysis

For analysis of Impα3/β binding to TDP-43-tev-MBP-His WT or NLS mutants, respectively, we performed binding studies using 5 µg of each purified protein in 300 µl PD-buffer (Transport Buffer (TPB): 20 mM Hepes pH 7.5, 110 mM KAc, 2 mM MgAc_2_, 1 mM EGTA, 2 mM DTT, 1 µg/ml each of aprotinin, leupeptin hemisulfate and pepstatin A supplemented with 2 mg/ml chicken ovalbumin) in absence or presence of 3.3 µM RanQ69L-GTP o/n at 4°C. Protein complexes were subsequently immobilized for 30 min at 4°C in total 400 µl PD-buffer by MBP-Trap beads (Chromotek), which had been pre-equilibrated in TPB supplemented with 20 mg/ml chicken ovalbumin. After washing 3x in 400 µl TPB lacking ovalbumin, bound proteins were eluted in 2x SDS sample buffer and analyzed by SDS-PAGE with subsequent Sypro-Ruby stain.

### Live cell imaging upon arsenite stress

To acquire time-lapse images of cytosolic condensation of the different EGFP-TDP-43 NLS-mutants, stable HeLa FlpIn EGFP-TDP-43 cell lines carrying the respective mutation (K82A, 83AAA or RK6A, respectively), were grown in ibiTreat 8well dishes (ibidi) and transfected with siRNA against endogenous TDP-43. EGFP-TDP-43 expression was induced using 20 ng/ml doxycycline. Imaging was performed in imaging media (DMEM Fluorobrite supplemented with 10% Tet-approved FBS) in an environmental chamber at 36.5°C at 5% CO_2._ Stress was induced by addition of Na-arsenite (final 0.5 mM; Sigma) directly to the imaging medium and images were acquired for 1h in 2.5 min intervals at high resolution using a Zeiss LSM900 equipped with Airy Detector (see below).

### Hormone inducible import assay (GCR_2_-EGFP_2_-reporter assay)

To analyze nuclear import of GCR_2_-EGFP_2_-tagged TDP-43 WT and NLS-mutants, nuclear import timelapse experiments were performed as previously described^67^. In brief, HeLa cells (gift by Marc-David Ruepp, KCL London) were grown for at least 2 passages in DMEM supplemented with 10% dialyzed FBS and subsequently transfected with the respective GCR_2_-EGFP_2_-TDP-43 constructs in ibiTreat 8-well dishes (ibidi) using Lipofectamine 2000. To reduce transfection stress, culture medium was exchanged 4-6h after transfection. Ca. 20-24h post-transfection, nuclear import was performed in DMEM Fluorobrite (Invitrogen) supplemented with 10% dialyzed FBS by addition of dexamethasone (final 5 µM, Sigma D4902) and time-lapse images acquired for 30 min in 2.5 min intervals in an environmental chamber set to 37°C and 5% CO_2_ using a spinning disc confocal (see below).

### SG association assay in semi-permeabilized cells

The analysis of TDP-43 SG association was performed as described^68^. In brief, HeLa cells were grown on high-precision (No. 1.5) poly-L-Lysine coated 12 mm coverslips, permeabilized using 0.005% digitonin in KPB (20 mM potassium phosphate pH 7.4, 5 mM MgAc_2_, 200 mM KAc, 1 mM EGTA, 2 mM DTT and 1 mg/ml each aprotinin, pepstatin A and leupeptin hemisulfate). After removing soluble components by several washes (4x 4 min in KPB on ice), nuclear pores were blocked by 15 min incubation in 200 µg/ ml wheat germ agglutinin on ice and cells were subsequently incubated with 100 nM recombinant TDP-43-tev-MBP-His_6_ in KPB at RT. Finally, cells were washed (3 x 5 min in KPB on ice) to remove unbound TDP-43-tev-MBP-His_6_ and processed by immunofluorescence using antibodies for G3BP1 (proteintech; #13057-2-AP) and MBP (proteintech, #66003-1-Ig) to visualize SGs and recombinant TDP-43-tev-MBP-His_6_, respectively. Samples were analyzed by confocal microscopy using a Zeiss LSM900 confocal microscope (see below).

### Immunofluorescence

Cells grown on coverslips were fixed for 7-10 min in 3.7% formaldehyde/PBS at RT, permeabilized for 5 min in 0.5% TX-100/PBS at RT and subsequently blocked in 5% donkey serum in PBS/0.1% Tween-20 (PBST) (all steps performed at RT). Cells were incubated with primary (for 1h at RT or o/n 4°C) and secondary antibodies (30-60 min at RT) diluted in blocking buffer. After 3 washes in PBST (5 min each), nuclei were counter stained for 5 min at RT using DAPI (0.5 µg/ml in PBS) and coverslips mounted with ProlongGlass Antifade (Life Technologies) and cured overnight at RT. For co-staining with EGFP-tagged TDP-43, AF647-labelled secondary antibodies were used to avoid spectral bleed-though/cross-talk. For the SG association assay, AF555- (for G3BP) in combination with AF488 (for TDP-43-tev-MBP-His_6_)-labelled antibodies were used to allow for visual discrimination of correctly permeabilized cells by G3BP1-staining. Subsequently, images of arsenite-induced cytosolic condensates of EGFP-TDP-43 were acquired by high-resolution imaging (Airy-Scan), images of the SG-association assay were acquired using by conventional confocal microscopy (see below).

### Solubility assay of EGFP-TDP-43 by RIPA extraction

Untreated or Na-arsenite treated HeLa cells (ca. 1×10^6^ cells each) were harvested by scraping into PBS with subsequent centrifugation at 1,200 g for 5 min. The cells were lysed using 200 µl RIPA buffer (50 mM Tris-HCL pH 8, 150 mM NaCl, 1% NP-40, 0.5% deoxycholate, 0.1% SDS) supplemented with 625 U benzonase (made by IMB protein production core facility) and 1x Sigma protease inhibitor mix. After 15 min incubation on ice, cells were sonicated for 45 sec using a BioRuptorPico (Diagenode) and 10% of the lysate retained as input. The rest of the lysate was centrifuged at 21,000 g for 30 min at 4°C to separate RIPA-soluble proteins from insoluble proteins. The pellet was once washed with RIPA buffer, followed by 45 sec sonication in the BioRuptorPico and centrifugation for 30 min at 21,000 g and 4°C. The final RIPA insoluble pellet was lysed by sonication (settings: 45 sec, high) in 40 µl Urea buffer (7 M Urea, 2 M Thiourea, 4% CHAPS, 30 mM Tris-HCl, pH8.5) using the BioRuptorPico. Samples were subsequently analyzed by western blotting using an EGFP-specific antibody (proteintech 66002-1-Ig) and band intensities determined by densitometry measurements (see below). Note, the 4.5x overrepresentation of the pellet fraction used for reasons of visibility (Figure 4) was corrected in the quantification.

### SEC analysis of purified TDP-43-tev-MBP-His_6_

To address if NLS-mutants of recombinant TDP-43-tev-MBP-His_6_ were dimerization-deficient, 50 µl of a 10 mg/ml TDP-43-tev-MBP-His_6_ were loaded onto a Superdex 200 Increase 10/300 GL column (GE-Healthcare) equilibrated in storage buffer (50 mM Tris pH8, 300 mM NaCl, 5% glycerol, 2 mM TCEP) and their elution profile was recorded at 280 nm. Every 10th A280 measurement point was subsequently plotted against the column volume using Graph Prism 9.

### Nuclear Magnetic Resonance (NMR) spectroscopy

^1^H-^15^N HSQC NMR spectra for TDP-43 NTF (aa 1-266) WT and RK6A were recorded at 298 K on an 800 MHz Bruker Avance NeoX NMR spectrometer equipped with a TCI cryoprobe. All samples were in 50 mM MES pH 6.1, 2 mM TCEP and 10% D_2_O was added for the lock. Characteristic and unambiguous chemical shifts of TDP43 NTFs (aa 1-266) were transferred using the deposited assignment of TDP43 N-terminal domain dimer and TDP43 RRM1-RRM2 (BMRB 30345 and 27613, respectively). Spectra were processed using TOPSPIN 5.0 (Bruker Biospin, Ettlingen, Germany) and analyzed using CcpNmr 2.5^69^.

### MD simulation

The condensates formed by full-length TDP-43 protein chains were simulated using the rescaled Martini 3 force field, with a protein–water interaction rescaling factor λ set to 1.06^70,71^. This rescaling was applied uniformly to all protein beads.

#### Single chain process

The atomistic structure of full-length TDP-43 was obtained from the AlphaFold Protein Structure Database (entry AF-Q13148-F1-v6)^72–74^. Coarse-graining was performed using the martinize2 tool within the vermouth 0.14.1 Python library^75^. Secondary structure assignment for structured regions was carried out using the STRIDE algorithm^76^ via the STRIDE Web interface^77^. Elastic networks were applied between beads with distances ranging from 0 to 0.85 nm, using a force constant of 700 kJ mol⁻¹ nm⁻² ^78^. These elastic bonds were specifically applied to structured regions of the protein, corresponding to residue ranges 4–76, 105–176, 192–259, and 320–332. Elastic restraints, as well as dihedral and angle potentials, were removed in unstructured regions. The modified Martini 3 force field was rescaled by scaling the Lennard–Jones (LJ) well depth ε using a factor λ = 1.06, applied uniformly to all protein – water interactions.

#### Chain assemblies

A total of 60 protein chains were simulated in a slab geometry with dimensions of 20 × 20 × 90 nm using the GROMACS 2024 software package^79,80^, with periodic boundary conditions applied in all directions. The simulation systems were neutralized by adding NaCl at a concentration of 150 mM. Lennard–Jones and Coulomb interactions were treated using the Verlet cutoff scheme, and electrostatics were handled with the reaction-field method. Production simulations were conducted under NPT conditions, with temperature maintained at 300 K using a velocity-rescaling thermostat^81^ and pressure controlled at 1 bar with the Parrinello–Rahman barostat^82^. A time step of 20 fs was used for integration.

To ensure statistical reliability, the WT condensates were simulated in three independent replicates, each initialized with a different spatial distribution of the protein chains. The simulation for the RK6A variant was performed using the same system parameters as for TDP-43 WT and corresponds to a single, representative simulation replicate.

### Densitometry measurements (Sypro-Ruby gels and Western blots)

Densitometry measurement of band intensities (I) of supernatant and pellet fractions for TDP-43 was performed in the corresponding detection software for Sypro-Ruby gels (Image Lab Software, Bio-Rad Laboratories) or western blots (Image Studio Software; Li-cor). To determine the relative solubility of individual proteins, the relative amount of the respective protein in the supernatant was calculated as the percentage of the sum of supernatant and pellet: % soluble = I_superatant_/ [I_supernatant_ + I_pellet_].

### Image acquisition and analysis

For display, all images were processed and assembled using Fiji/Image J applying linear enhancement for brightness and contrast. Unless noted differently, equal processing was applied for images of the same channel within the same experiment. All graphs and statistical analyses were performed using Graph Prism 9 (details indicated in the respective figure legends).

#### Phase contrast imaging of TDP-43 condensates

All images were acquired in an automated fashion on an inverted Zeiss Observer 7 equipped with Definite Focus, a 63x/1.4 Oil Ph3 objective and an AxioCam 705 Mono camera using the Zeiss ZEN software. If applicable, phase contrast images were combined with structural illumination (Zeiss Apotome) for images of fluorescent RNA using a Colibri 5 light source equipped with a 630 nm LED and an emission bandpass filter (681/45).

#### Confocal microscopy

Images for the SG -association assay in semi-permeabilized cells were acquired on a Zeiss LSM900 using a 63x/1.4 oil objective with sequential unidirectional scanning at 71 nm pixel size at 16 bit and two-fold frame averaging. The following fluorescence and detection settings were used: AF555 (561nm laser): 540-700 nm; AF488 (488 nm laser): 410-540 nm; DAPI (405 nm laser): 400-600 nm. Images were analyzed in an automated fashion using machine - learning driven segmentation (Zeiss Intellesis) to measure the AF488-signal (i.e., TDP-43-tev-MBP-His_6_) within SGs (defined by the G3BP1-channel). All values were background subtracted and normalized to TDP-43 WT.

#### High resolution confocal microscopy (AiryScan)

High resolution images were acquired on a Zeiss LSM900 equipped with AiryScan2 Detector. Timelapse images were acquired with a Plan-Apo 40x/1.2 water auto-immersion objective at 50 nm pixel size (SR-2Y multiplex mode) with bidirectional scanning using the 488 nm laser and a detection window of 490-620 nm. Image analysis was performed using an in-house developed jupyter notebook python script to quantify cellular condensate number and cluster formation. In brief, individual cells were marked as ROI in Napari viewer^83^ and ROIs expanded over time. Condensates within ROIs were detected as local maxima by the skimage library blob-log algorithm using equal thresholds within the same replicate to optimally detect cytosolic condensates. Subsequently, cluster formation was analyzed by sklearn DBSCAN algorithm. Script development was supported in parts by use of AI.

For high resolution images for cytosolic EGFP-TDP-43 condensates, images were acquired at 16-bit with a 63x/1.4 oil immersion objective at 35 nm pixel size (SR mode) by sequential unidirectional scanning and two-fold frame averaging. The following fluorescence settings were used: AF647 (640 nm laser): 450-700 nm; EGFP (488 nm laser): 490-620 nm; DAPI (405 nm laser): 400-495 nm.

#### Spinning disc confocal microscopy

Imaging of nuclear import rates was performed at the central microscopy core facility (CMCF) of the IMB with an inverted spinning disc confocal (Visiscope 5 Elements, Visitron) built on a Nikon Ti2 stand equipped with a confocal spinning disc (CSU-W1, Yokogawa, Tokyo, Japan) equipped with a 488 nm laser line for excitation, a 565/133 bandpass emission filter (Semrock) and a 50 µm pinhole diameter. Images were acquired using a 60x/1.2 NA water immersion objective and a sCMOS camera (Prime BSI, Photometrics) with full-frame (2048 x 2048 pixels). Image analysis was performed using Fiji/ Image J (1.54). If applicable, xy drift of cells was compensated by the StackReg plugin before downstream analysis. To determine the nuclear import rate, at least 40 cells per replicate were analyzed by measuring a respective circular ROI (region of interest) in nucleus (N) and cytoplasm (C), respectively and plotting them after background subtraction as N/C ratio over time.

## Results

### Phase separation of TDP-43 requires basic amino acids in its disordered NLS-region

Different computational tools predict that not only the C-terminal LCD but other regions of TDP-43 have a high degree of disorder, in particular the N-terminal region comprising the NLS (aa 79-101; Fig. 1A). To experimentally test if the disordered NLS-region is involved in TDP-43 phase separation, we purified recombinant TDP-43 either as wildtype (WT) protein or with the entire NLS-region deleted (delNLS) or replaced with a disordered uncharged linker (NLSrepl) from bacteria and compared their condensate formation propensities after Tev protease-mediated removal of the C-terminal MBP-solubility tag (Fig. 1B-G, Fig. S1A-C). At 5 µM protein concentration, TDP-43 WT readily formed spherical condensates, which increased in size at higher protein concentrations. In stark contrast, TDP-43 delNLS did not form any visible condensates up to 40 µM, indicating a crucial role of the disordered NLS-region for TDP-43 phase separation. Interestingly, replacing the NLS-region with a disordered, uncharged linker sequence (NLSrepl) restored condensate formation, albeit only at higher concentrations (10 µM) and with different morphology than WT (larger condensates exhibiting some surface wetting at higher protein concentrations) (Fig. S1C). This indicates that not only the disordered nature of the NLS-region but also its sequence context governs TDP-43 phase separation.

**Figure 1:**
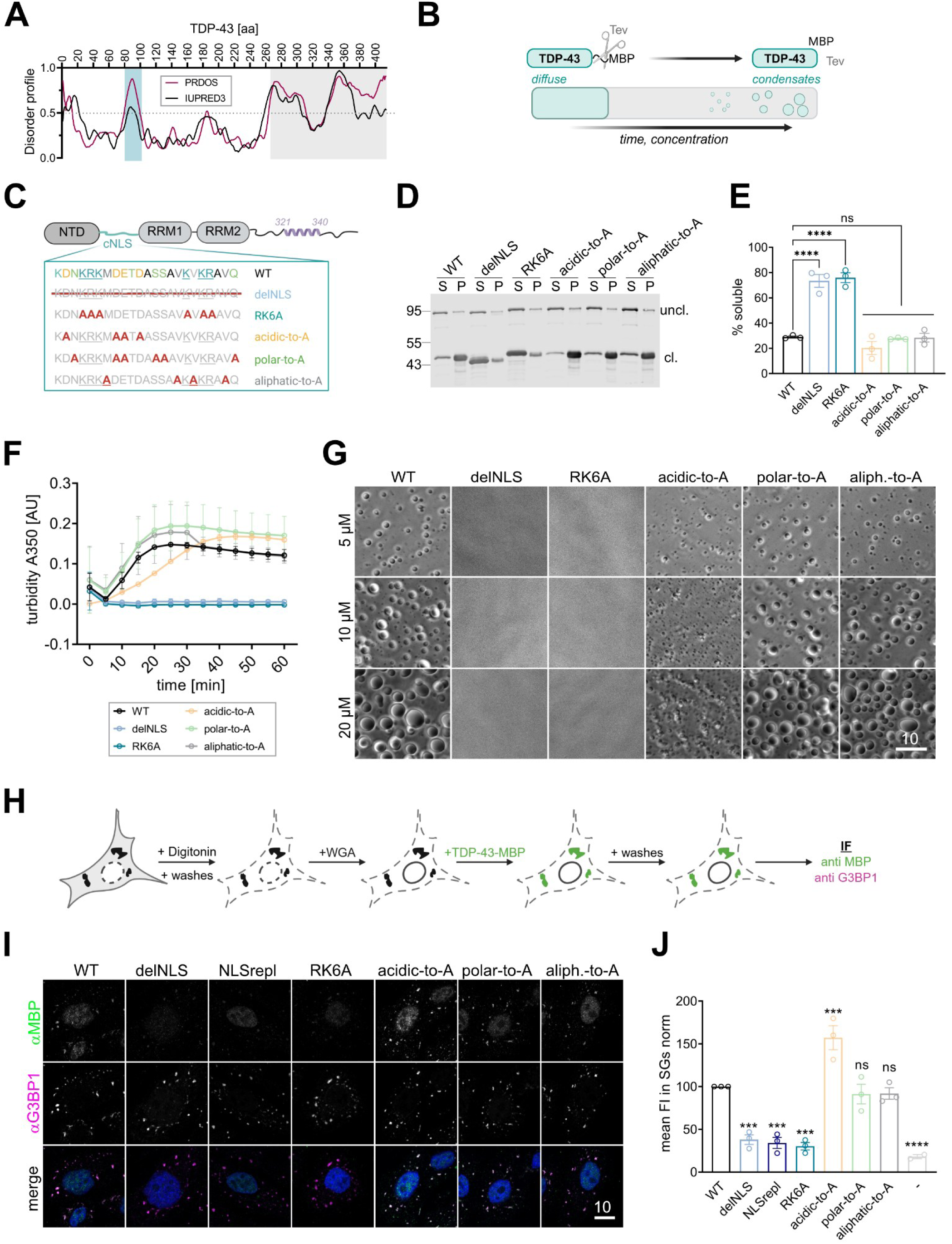
Basic residues in the disordered cNLS-region are required for phase separation of TDP-43. A) Disorder prediction for TDP-43 (using IUPRED3 and PrDOS ^64,65^) with region corresponding to NLS-region and C-terminal LCD highlighted by the turquoise and grey area, respectively. B) Tev-induced cleavage of the MBP-solubility tag induces TDP-43 phase separation in a time and concentration-dependent manner. C) Cartoon showing domain structure of TDP-43 (NTD, N-terminal domain, RRM, RNA-binding motif). The cNLS sequence with the respective amino acid classes (basic: turquoise; acidic: orange, polar: green; aliphatic: grey) are highlighted with the bipartite cNLS motif underlined. Systematic mutations of NLS-region are indicated below. D) Sedimentation assay to quantify condensation of TDP-43 WT and indicated NLS-mutants at 5 µM detected by Western blot (N-terminal TDP-43 antibody). Equal fractions of supernatant (S) and pellet (P) are shown. E) Quantification of percentage of soluble, cleaved TDP-43 calculated as [S/(S+P)] for 3 independent replicates. ****<0.0001 by One-way Anova with Dunnett’s multiple comparison test to WT. ns, non-significant. F) Turbidity assay (absorbance at 350 nm) of phase separation for TDP-43 and indicated NLS mutants over time (mean ± SD for two independent replicates). G) Representative phase contrast images of condensates for TDP-43 WT and indicated NLS mutants across 3 concentrations. Bar, 10 µm. H) Cartoon demonstrating general principle of the SG association assay in semi-permeabilized cells shown in (I- J). (I) representative confocal fluorescence microscopy images of a SG association assay in semi-permeabilized HeLa cells for indicated TDP-43-tev-MBP-His_6_ variants. (J) Quantification of the respective SG association of the TDP-43 variants normalized to WT, mean of three independent experiments ± SEM; ns, not significant, *** <0.0002, **** <0.0001 by 1-way ANOVA with Dunett’s multiple comparison test compared to WT. (-) indicates background signal in absence of recombinant TDP-43.

We therefore created several TDP-43 variants in which we systematically mutated the different amino acid classes (basic, acidic, polar and aliphatic) in the disordered NLS-region to alanine to analyze the relevance of the different amino acid types for TDP-43 condensation (Fig. 1C and S1A) in different, complementary phase separation assays. First, we quantified the level of TDP-43 solubility as a direct measure for its condensation behavior in a sedimentation assay. As shown in Figure 1D (see Fig. 1E for quantification), TDP-43 WT is predominantly found in the pellet fraction (∼30% solubility), similar to the variants where polar or aliphatic NLS residues were mutated to alanine. Mutation of acidic residues resulted in a small, but non-significant decrease in TDP-43 solubility. Interestingly, mutation of all basic amino acids (RK6A) in the NLS-region caused a strong increase of soluble TDP-43, similar to what is observed when the entire NLS-region is deleted (∼70% solubility). Next, we measured TDP-43 condensation over time in a plate reader turbidity assay. Here, the time-dependent increase in turbidity for TDP-43 mutants carrying mutations in polar or aliphatic NLS-residues was comparable to TDP-43 WT, while TDP-43 with mutations in acidic NLS residues exhibited delayed onset of turbidity. In contrast, both the delNLS as well as the RK6A variant did not show any turbidity over the time course of 1 hour (Fig. 1F). Finally, we utilized phase contrast microscopy to compare the morphology of TDP-43 condensates at different protein concentrations (Fig 1G). In line with our results from sedimentation and turbidity assays, deletion of the NLS or mutation of basic amino acids in the NLS (RK6A) completely prevented formation of condensates across all examined protein concentrations (5-20 µM). Mutation of acidic residues resulted in small, more amorphous condensates, possibly explaining the observed delayed increase in turbidity. Mutation of polar or aliphatic resides had no influence on morphology or size of TDP-43 condensates.

As previous studies had described a stimulation of TDP-43 phase separation by addition of specific TDP-43 target RNAs^84^, we also tested the influence of such an RNA sequence on TDP-43 RK6A condensation. However, TDP-43 RK6A phase separation could not be stimulated by addition of a known TDP-43 target RNA (Fig. S1D), even though the RNA was able to partition into TDP-43 WT condensates at substoichiometric concentrations and at equimolar concentrations was able to suppress TDP-43 WT condensates. Importantly, this lack of stimulation is not due to reduced RNA-binding capacity of the delNLS and RK6A variants, as both bind RNA similarly to TDP-43 WT as determined by EMSA (Fig. S1E-G). TDP-43 is a known component of cytoplasmic stress granules (SGs)^15,19^ and we have previously shown that partitioning of recombinant TDP-43 into preformed SGs in semi-permeabilized cells correlates with its phase separation propensity^35^. When we quantitatively compared the SG association of our various TDP-43 NLS variants (Fig. 1H-J), mutations suppressing TDP-43 phase separation (delNLS, NLSrepl and RK6A) all caused a strong decrease in SG partitioning compared to WT in accordance with our condensation experiments (Fig. 1I, see J for quantification). In contrast, mutation of acidic residues resulted in enhanced SG association, while mutation of polar and aliphatic residues had no effect. In summary, our data show that not only the C-terminal LCD is important for TDP-43 phase separation, but also the disordered NLS-region, in particular the basic amino acids.

#### Basic amino acids in the disordered NLS-region engage in multiple contacts across TDP-43

The NLS of TDP-43 contains two arginines and four lysines (see Fig. 1C). Arginines are often stronger drivers of phase separation than lysines^85–87^, although a role of lysines in complex coacervation with RNA and phase separation of the Tau protein has been reported^88^. To analyze the relevance of arginines and lysines in the NLS motif in TDP-43 phase separation, we mutated them separately to alanine (R2A, K4A; see Fig. 2A and S2A) and studied condensate formation of the respective recombinant proteins using phase contrast microscopy (Fig. 2B). While mutation of the two arginine residues (R2A) had only a minor influence on TDP-43 condensation, mutation of four lysines in the NLS-motif (K4A) suppressed phase separation at 5 µM concentration. Both R2A and K4A mutant showed formation of enlarged condensates at high protein concentrations likely due to fusion events. The disordered NLS-region (aa 79-101) contains an additional lysine (K79), which is not part of the NLS motif (Fig. 2A). Interestingly, upon mutating all five lysines to alanines (K5A), phase separation of TDP-43 was as strongly suppressed as for the RK6A mutant, pointing to an important role of these lysine residues (Fig. 2B). To test if suppression of phase separation could be caused by the decreased net charge per residue (NPCR) of the K5A mutant (NCPR: -0.022) compared to TDP-43 WT (NCPR: −0.010), we included a different TDP-43 variant with comparable NCPR (S5D; NCPR: −0,022)^35^ in our analysis (Fig. 2C,D). However, as described previously^35^, the onset of phase separation of this mutant was similar to WT (Fig. 2D), the main difference being larger condensates at higher concentrations and a pronounced wetting effect, indicating increased liquidity. As K79 is directly proximal to the multimerization domain of TDP-43, we cannot exclude an effect of the K79A mutation on TDP-43 dimerization. The K79A mutation alone showed no diminished condensation properties upon Tev protease cleavage when analyzed by phase contrast microscopy (Fig. S2C), suggesting no major effect of the K79A mutation on TDP-43 self-interaction. In contrast, the well-known dimerization-deficient 6M mutant^6,46^ was completely phase separation deficient (Fig. S2B,C). To assess whether the condensation deficiency of some of our TDP-43 NLS variants was caused by reduced dimer formation, we analyzed them as MBP-fusion proteins by analytical size exclusion chromatography (SEC) in comparison to WT (dimerization-competent) and 6M (monomeric)^6,46^. Interestingly, the TDP-43 delNLS and TDP-43 K5A fusion protein showed an elution profile more similar to the monomeric 6M mutant, while the RK6A protein behaved similar to WT (Fig. S2D). This indicates that at least parts of the disordered linker domain between NTD and RRM1 might be involved in dimer formation and/or stabilization, yet the RK6A mutant showed no obvious defect in TDP-43 dimerization when fused to the MBP-solubility tag at slightly basic pH.

**Figure 2:**
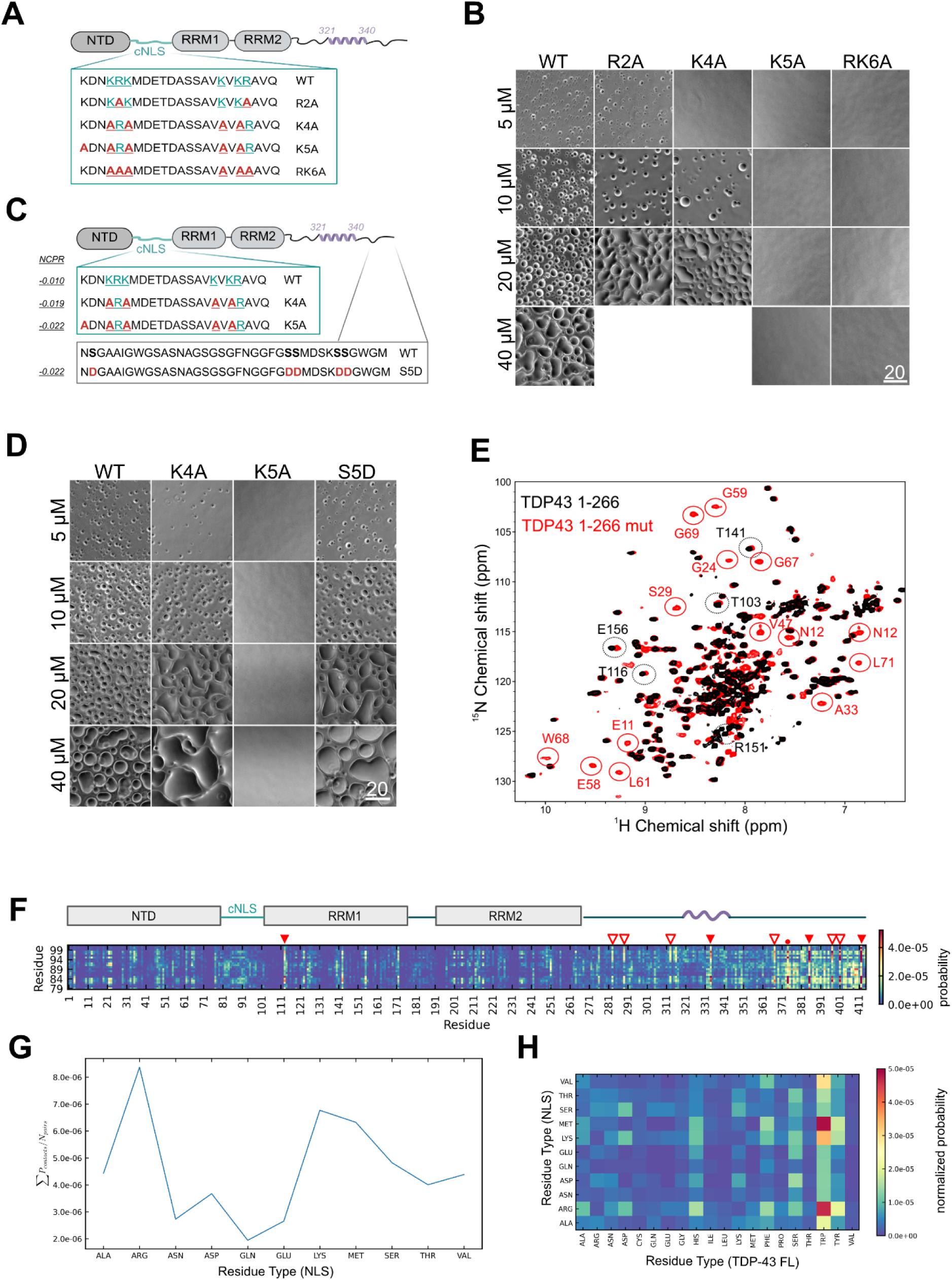
TDP-43 NLS is involved in a multiple inter- and intramolecular interactions. A, B) Cartoon (A) and representative phase contrast microscopy images for phase separation behavior (B) of TDP-43 WT and mutants in which either lysines (K) of arginines (R) of the disordered NLS-region have been mutated to alanine (A). C, D) Cartoon (C) for TDP-43 WT and NLS-lysines mutants analyzed by phase contrast microscopy in (D) compared to a TDP-43 mutant carrying indicated serine-to-aspartic acid mutations in the LCD (S5D) with similar net charge per residues (NCPR; calculated by CIDER^63^). Bar, 20 µm. E) Overlay of ^1^H,^15^N HSQC spectra of uniformly ^15^N-labeled TDP43 NTF WT (black) and NTF RK6A (red) at 100 µM. Chemical shift perturbations in the RRM1 are indicated by black circles, while signal recovery from the RK6A mutant in the NTD are encircles red. F) Inter-chain contact map from adapted Martini3 elastic network coarse-grained model of the disordered TDP-43 NLS-region (aa 79-101) over the full-length sequence. The reference TDP-43 domain structure (cartoon above) and probable interaction sites with either tryptophan (full arrowheads; RRM1: W113; LCD: W334, W385, W412) or phenylalanine (empty arrow heads; LCD: F276, F283, F289, F313, F367, F397, F401) and tyrosine (circle; Y373 in LCD) are indicated. G) Contact frequency of residue types within the disordered NLS-region against residues in full-length protein chains normalized by the respective sum of residue pairs. H) Residue type inter-chain contact map of disordered NLS-region over full-length TDP-43, contact probability normalized by number of interaction pairs.

Since a recent publication demonstrated stacking of the TDP-43 NTD including the NLS-region (aa 1-101) onto the RRM domains^89^, we performed NMR spectroscopy to investigate whether the basic amino acids in the disordered NLS-region are involved in this intramolecular interaction. Comparison of ^1^H-^15^N HSQC spectra of TDP-43 N-terminal fragments (NTF, aa 1-266) that had either the WT sequence or harbored the RK6A mutation, exhibited chemical shift perturbation of ^1^H-^15^N cross peaks that could be assigned to the RRM1 domain, indicating a change in the local environment of RRM1 for the RK6A mutant (Fig. 2E; black circles). Additionally, the RK6A mutant NTF also showed signal recovery of peaks corresponding to the N-terminal dimerization domain (NTD) (Fig. 2E; red circles). Given that these signals were absent in the WT NMR spectra, which could be due to line broadening caused either by conformational exchange of the NTD or interaction of the NTD with other parts of TDP-43, the recovery of signals in the RK6A mutant is indicative of reduced destabilization or self-interactions of the NTD and/or the NLS itself.

To further support the evidence that the basic amino acids in the NLS-region are involved in TDP-43 self-interactions, we turned to molecular dynamics (MD) simulations. We used a recently developed near-atomic-resolution, adapted Martini3 elastic network coarse-grained model to simulate full-length TDP-43 phase separation, which indicates inter-domain contacts of the LCD with other regions of TDP-43 and highlights the importance of aromatic residues, such as phenylalanine or tryptophan, in the C-terminal LCD for TDP-43 condensation^48^. Interestingly, these simulations demonstrated that also the disordered NLS-region (aa 79-101) has multiple intermolecular contact sites across full-length TDP-43, with local high probability interaction sites especially in the C-terminal LCD, and to some extent also in the RRM1 and NTD, or weakly in the NLS-region (Fig. 2F). In line with our *in vitro* experiments, these interactions were predominantly mediated by arginine and lysine residues in the disordered NLS-region (see Fig. 2G for contact probabilities normalized by residue pair abundance; see Fig. S2E for non-normalized data). They show high contact probability with several aromatic residues in the RRM1 and LCD (Fig. 2F), from which in particular tryptophan shows high potential as interaction sites (Fig. 2H for contact map normalized by residue pair abundance; see Fig. S2F for non-normalized contact map). The close proximity to arginine and lysine within the NLS is likely also the reason for the high contact probability for methionine (aa 85; Fig. 2H) rather than reflecting a distinct interaction preference. To verify the role of basic amino residues in the NLS, we also performed simulations for TDP-43 RK6A using the same parameters as for TDP-43 WT. Indeed, inter-chain contacts were reduced for this mutant, as demonstrated by reduced contacts of the corresponding residues in the NLS region as well as key residues in the LCD, NTD and RRM (Fig. S2G).

In summary, our data show a crucial role of the positively charged amino acids in the disordered NLS-region for TDP-43 phase separation, as these residues mediate multiple inter-domain interactions, in particular with aromatic residues in the C-terminal LCD, but also some interactions with the RRM1 and NTD domain.

#### Basic residues in the NLS, NTD dimerization and C-terminal aromatic residues are equally important for TDP-43 condensate and cluster formation

To further examine the relevance of the basic residues in the NLS, we compared their impact on TDP-43 phase separation with mutations reported to interfere with TDP-43 self-assembly, or mutated their main interacting residues in the LCD as identified in our MD simulation. To this end, we purified additional TDP-43 variants carrying mutations in all three tryptophans in the LCD (W3A; W334/385/W412A), in six phenylalanines that show high contact probability in our MD simulation (F6A; F276/283/289/367/397/401), leaving the phenylalanine close to the CR intact, as to not compromise α-helix formation^90^, or made a combination of both mutations (F6W3A) (Fig. 3A, S3A). Additionally, we examined a dimerization-deficient TDP-43 mutant (6M)^6^, which previously was shown to suppress TDP-43 condensation *in vitro*^46^ (see also Fig. S2C), and a variant lacking the residues involved in α-helix formation in the CR (aa 321-340, delCR) (Fig. 3A, S3A). Deletion of the CR previously has been reported to interfere with phase separation of the C-terminal LCD of TDP-43^38^ as well as nuclear condensation of TDP-43 in _cells43,44,91._

**Figure 3:**
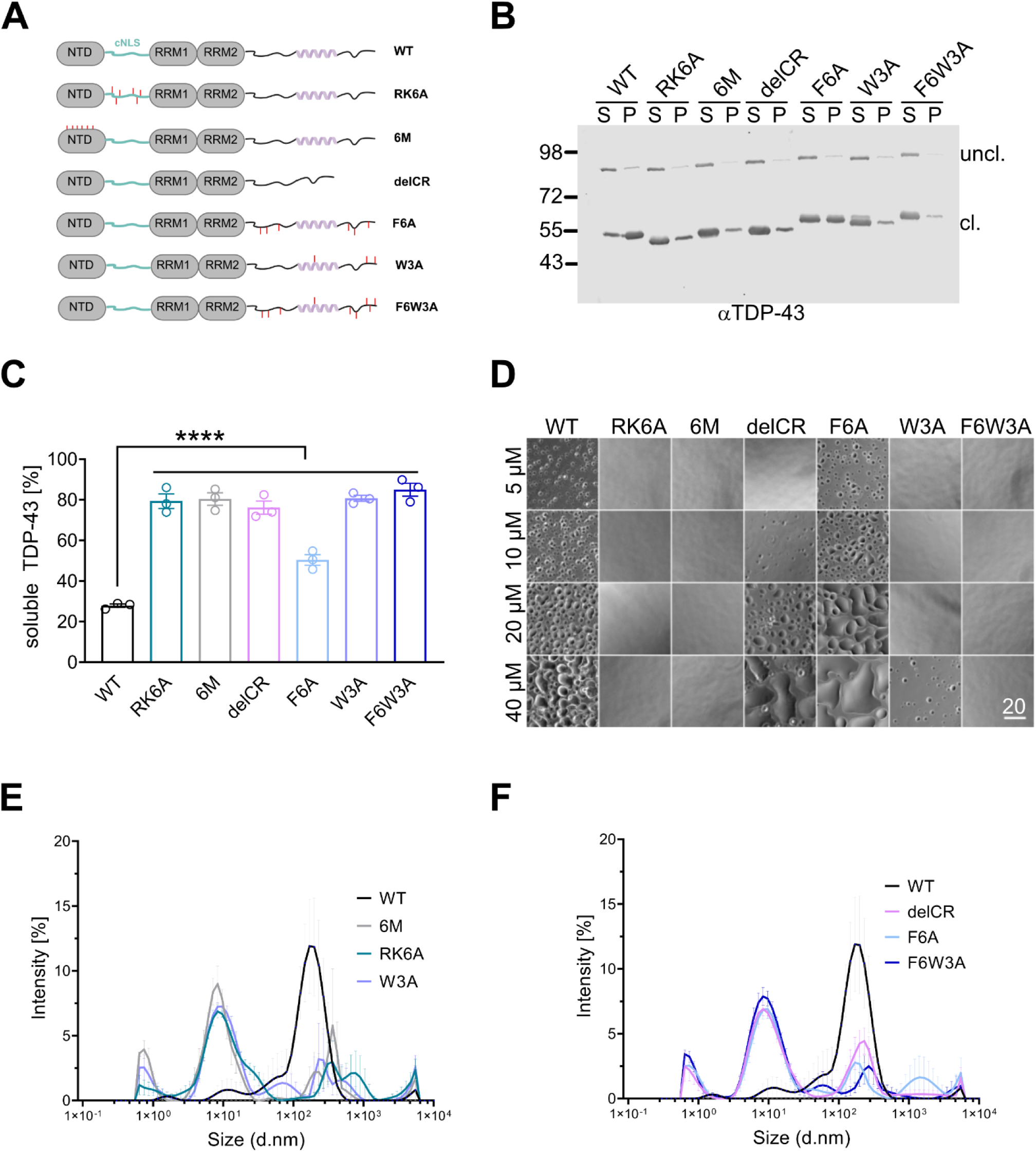
Dimerization, TDP-43 NLS and C-terminal aromatic residues are important for micron-sized condensate as well as nanocluster formation of TDP-43. A) Cartoon depicting mutants of recombinant TDP-43 used to compare phase separation propensity of TDP-43 WT and RK6A with various mutants of TDP-43 known or predicted to show reduced condensation behavior. (B-F) Analysis of phase separation behavior by either sedimentation analysis (B, C), phase contrast microscopy (D) or DLS (E, F) 1h after Tev protease cleavage. C) Quantification of three independent sedimentation assays using a N-terminal TDP-43 antibody as shown in (B), mean ± SEM of 3 independent experiments, **** < 0.0001 by 1-way ANOVA with Dunetts multiple comparison test to WT. E, F) Intensity profile of nanocluster formation as determined by DLS for indicated TDP-43 variants at subcritical concentrations (0.7 µM), mean ± SEM. All mutants exhibit reduced self-assembly indicated by the appearance of smaller species. Note, WT is duplicated in E and F for better visibility.

When we analyzed the level of soluble TDP-43 upon Tev protease cleavage of the MBP-solubility tag in a sedimentation assay, nearly all mutant variants showed strongly enhanced solubility compared to the WT protein (Fig. 3B, see C for quantification), indicating reduced self-assembly. Alone the F6A mutant showed an intermediate solubility (∼50%). Visual inspection of condensate formation by phase contrast microscopy at varying concentrations in a physiological buffer, revealed interesting differences in the condensation behavior of the different variants: Surprisingly, deletion of the CR region suppressed phase separation only at low protein concentration (5 µM), but resulted in the formation of few, small condensates at intermediate concentration (10 µM) and large, liquid-like condensates with a strong wetting effect at high concentration (40 µM). In line with our sedimentation analysis, the F6A mutant showed no significant deficiency in condensate formation, yet exhibited a strong droplet fusion and wetting phenotype at high concentrations (20-40 µM). Mutation of the three tryptophans (W3A) strongly suppressed condensate formation across all concentrations, with small condensates appearing only at 40 µM. Additional mutation of phenylalanines (F6W3A) resulted in even stronger suppression of condensate formation, indicating a weak contribution of the phenylalanines on TDP-43 phase separation. Notably, in addition to the F6W3A mutant, only the dimerization-deficient 6M mutant was as deficient in TDP-43 phase separation as the NLS RK6A variant, all three variants lacked visible condensates even at 40 µM (Fig. 3D).

To test whether our recombinant TDP-43 can also form nano- or mesoscale assemblies at subsaturated concentrations as recently reported for FUS and other RBPs^50,51^ and whether this nano-/mesoscale self-assembly is also influenced by mutations that suppress visbleTDP-43 condensate formation, we applied dynamic light scattering (DLS) in physiological buffer conditions at 0.7 µM 1h after Tev protease cleavage. When displayed as scattering intensities against the apparent hydrodynamic radius (d.nm.), a dominant peak at about ∼100-300 nm was seen for TDP-43 WT (Fig. 3E, F; note that the data for TDP-43 WT are duplicated in both panels). As the monomeric form of TDP-43 is expected to have a size of ∼ 4.6 nm (or ∼9 nm for a dimer), the observed signal indicates the formation of mesoscale assemblies by TDP-43 WT. In contrast, all mutants exhibited strongly reduced scattering (see Fig. S3B for autocorrelation function) and smaller species appeared in the intensity profile (Fig. 3E, F), suggesting that all tested mutations reduce TDP-43 self-assembly at sub-critical concentration.

Together, our data show that the basic amino acids in the disordered NLS-region of TDP-43 are crucial for its self-assembly and phase separation, both in the range of visible condensates and mesoscale cluster regime.

#### Mutations abolishing TDP-43 nuclear import show a gradual deficiency in TDP-43 self-assembly in vitro

Mutations of basic amino acids in the NLS motif interfere with import of TDP-43 in the nucleus and are therefore often used to mimic disease-linked cytosolic mislocalization in cell or animal models aimed at recapitulating TDP-43 proteinopathy. We next set out to compare different mutations commonly used in the field to assess their impact on TDP-43 self-assembly. The RK6A mutant carries alanine substitutions of all six basic residues in both parts of the bipartite NLS and has previously been used in cellular studies^92–94^. Other studies employed mutations in the first half of the bipartite cNLS, for example the commonly used rNLS TDP-43 mouse model^95^ and other cellular studies^24,29,96–98^. It is usually referred to as ΔNLS (defective NLS) and carries alanine substitutions in the KRK motif (aa 82-84). To avoid confusion with our construct harboring a complete NLS deletion (delNLS), we will from here on refer to it as ‘83AAA’ (Fig. 4A). More recently, the first half of the cNLS of TDP-43 was shown to serve as the main binding site for TDP-43’s cognate nuclear import receptor Impα/β^14^. Here, in particular, lysine 82 plays a critical role in mediating binding to Impα and consequently, mutation of K82 is sufficient for abolishing binding to Impα/β and causing cytosolic mislocalization of TDP-43^14,99,100^.

**Figure 4:**
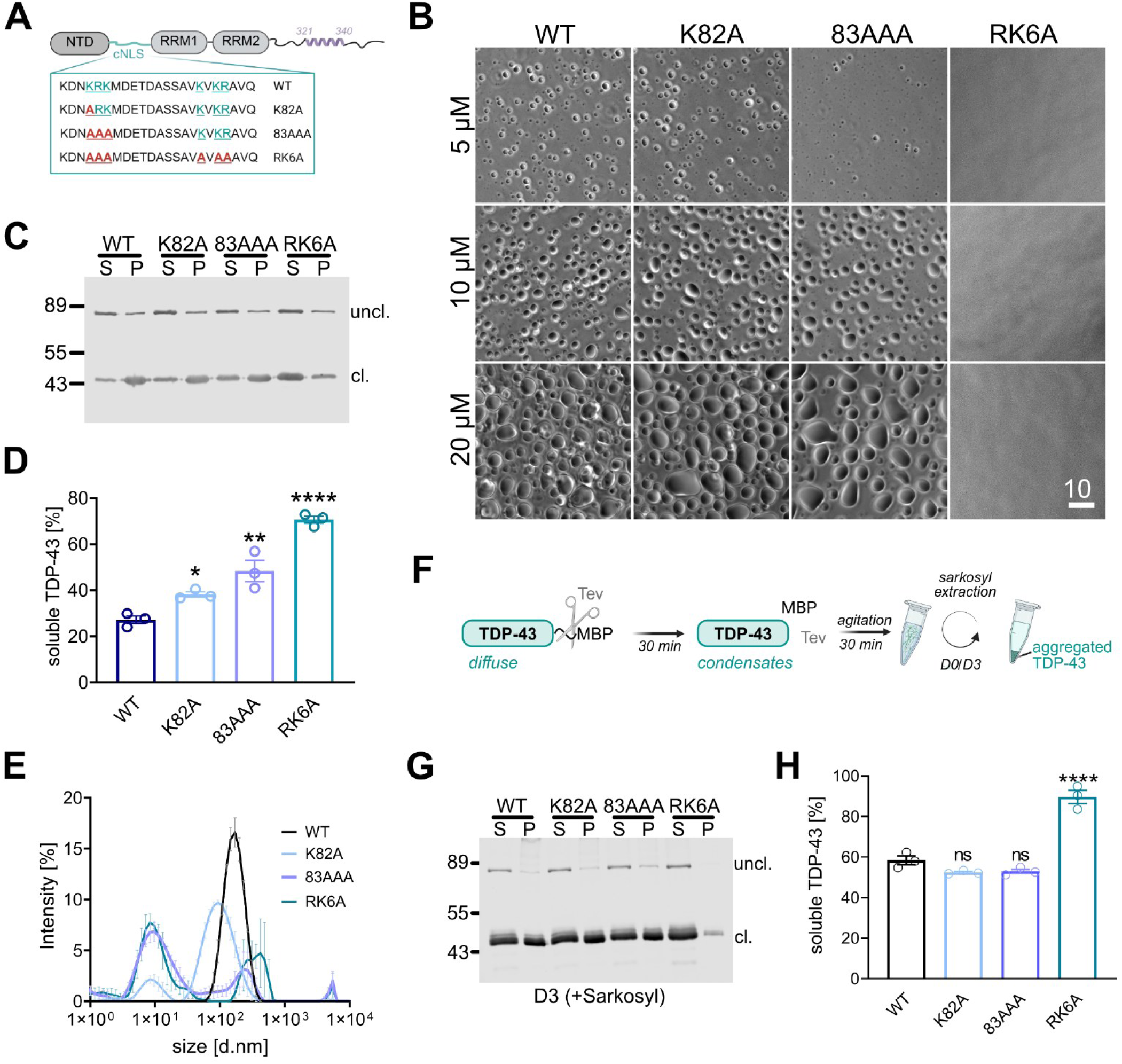
Differential phase separation and aggregation behavior of commonly used TDP-43 import-deficient mutants in vitro. A) Cartoon depicting mutants of recombinant TDP-43 carrying commonly used cNLS mutations to abolish nuclear import. B) Representative phase contrast images of TDP-43 nuclear import-deficient mutants shown in (A). C, D) Sedimentation assay to quantify condensation of TDP-43 and indicated NLS-mutants at 5 µM detected by Western blot (C) (N-terminal TDP-43 antibody). Equal fractions of supernatant (S) and pellet (P) are shown. D) Quantification of percentage of soluble cleaved TDP-43 calculated as [S/(S+P)] for 3 independent replicates. * < 0.0332, ** < 0.0021, **** < 0.0001 by One-way Anova with Dunett’s multiple comparison test to WT. E) DLS measurements displayed as intensity profile for import-deficient TDP-43 variants at subcritical concentrations (0.7 µM), shown as mean ± SEM. Note that the single mutant K82A to some extent, but in particular the 83AAA and RK6A mutant exhibit reduced self-assembly as indicated by the appearance of smaller species. F) Scheme for aggregation assays shown in (G-H). Phase separation of recombinant TDP43-tev-MBP-His_6_ was induced by Tev-mediated cleavage for 30 min before shaking for 30 min at RT to induce aggregation. (G, H) Representative Western blot (G) of sedimentation analysis after sarkosyl extraction. Equal fractions of soluble (S) or insoluble (pellet, P) fractions were analyzed by Western Blot using a N-terminal TDP-43 antibody. (H) Quantification of percentage of sarkosyl-soluble, cleaved TDP-43 calculated as [S/(S+P)] for 3 independent replicates at day 3 (D3), mean ± SEM by 1-way Anova with Dunnett’s multiple comparison test to WT.

We first verified that all these mutations in the cNLS of TDP-43 indeed interfere with binding to Impα/β by performing pulldown experiments with recombinant MBP-tagged TDP-43 WT, K82A, 83AAA or RK6A as a bait, in conjunction with recombinant Impα3 and Impβ in the absence or presence of RanQ69L-GTP (Fig. S4A). Addition of the GTPase-deficient RanQ69L in its GTP-bound form serves as a specificity control for the interaction, as it dissociates import complexes, thus mimicking the nuclear environment. Indeed, we detected specific binding of Impα3 and Impβ to TDP-43 WT in the absence but not presence of RanQ69L-GTP. As expected, none of the NLS mutants showed significant binding to Impα or Impβ, irrespective of the presence of RanQ69L-GTP. Next, we verified nuclear import deficiency of the different NLS mutants by live cell imaging. Previous studies reported varying levels of cytoplasmic mislocalization of TDP-43 mutated in K82 in a steady-state analysis^99,100^. We now employed our EGFP-tagged hormone-inducible import system, which traps EGFP_2_-tagged TDP-43 in the cytoplasm due to its fusion to two moieties of the hormone- binding domain of the glucocorticoid receptor (GCR)^35,61,67^. After addition of a steroid hormone (dexamethasone), this reporter is readily imported into the nucleus dependent on TDP-43’s intrinsic NLS and its import rates can be quantitatively determined by live cell imaging (Fig. S4B). In line with our pulldown experiments, the TDP-43 K82A mutant was as import-deficient as the more extensive RK6A mutant (Fig. S4C, see D for quantification), verifying previous steady state data^99,100^.

We then set out to characterize the phase separation behavior of these three NLS mutant TDP-43 variants both by visual inspection using phase contrast microscopy and sedimentation analysis. Interestingly, condensate formation showed a direct correlation to the number of mutations: In contrast to the strongly phase separation-deficient RK6A mutant, the 83AAA mutation showed reduced condensation at low concentrations (5 µM), while the single point mutant (K82A) showed condensate formation comparable to WT (Fig. 4B). In line with this, TDP-43 solubility as determined by sedimentation analysis also correlated with number of basic amino acid substitutions in the NLS (Fig. 4C, see D for quantification). As the RK6A mutant was strongly deficient in cluster formation at subcritical concentrations (Fig. 3E), we also subjected the K82A and 83AAA mutant to DLS measurements. Here, already the single mutation (K82A) resulted in the appearance of smaller species compared to WT (Fig. 4E, see Fig. S4E for autocorrelation curve indicating decreased scattering). The 83AAA mutant had a similar profile as the RK6A mutant, indicating a strong deficiency in mesoscale cluster formation. Lastly, we addressed the aggregation behavior of recombinant TDP-43 NLS-mutants. Resistance to detergents, such as sarkosyl, is a feature of pathological TDP-43 inclusions^101–104^. Therefore, we first used an established protocol to generate large, amorphous TDP-43 aggregates induced by short shaking after Tev protease cleavage in an aggregation-promoting buffer^35,66^ and after incubation for up to three days performed sarkosyl extraction and quantitative western blot analysis of the soluble and insoluble fractions (Fig. 4F). Directly after shaking (day 0), TDP-43 WT and all NLS mutants were nearly completely sarkosyl-soluble (Fig. S4F, see G for quantification). However, after three days of incubation at room temperature, TDP-43 WT showed strongly reduced solubility upon sarkosyl extraction (Fig. 4G; see H for quantification). Both the K82A and 83AAA mutant showed similar reduction in sarkosyl-solubility, indicating that they have a similar aggregation behavior as the WT protein. In stark contrast, the RK6A mutant was nearly completely solubilized by sarkosyl under those conditions, indicating decreased aggregation properties. In summary, our findings demonstrate a strong influence of the number of basic residue mutations in the NLS on TDP-43’s self-assembly behavior across length scales.

#### Level of cytosolic TDP-43 condensation is governed by the extent of NLS mutations

To transfer these *in vitro* findings into cells, we first quantified the level of SG association of the three different NLS mutants in our semi-permeabilized cell assay (Fig. S5A, see B for quantification). In line with our *in vitro* data, we found a gradual reduction in SG association of TDP-43 with increasing number of basic amino acid substitutions: The K82A mutant showed no significant change compared to WT, whereas the 83AAA mutant displayed significantly reduced SG partitioning, yet not as pronounced as the RK6A mutant (Fig. S5A,B), indicating that the number of basic amino acids in the NLS play an important role in SG recruitment of TDP-43. Next, we characterized the condensation behavior of these mutants in intact cells. To avoid overexpression artefacts and varying expression levels that can influence TDP-43 condensation, we generated stable HeLa cell lines using the established Flp-In T-REx system that enables stable integration of a single copy of a gene-of-interest under a dox-inducible promoter^105,106^. Western blot analysis confirmed near equal expression levels of EGFP-TDP-43 WT for all three NLS mutants, with the 83AAA and RK6A line showing modestly elevated levels of EGFP-TDP-43 compared to the WT and K82A line (Fig. S5C). We used a highly efficient TDP-43-specific siRNA to deplete endogenous TDP-43 from all stable lines, to avoid any influence of endogenous, condensation-prone TDP-43 in our analyses (Fig. S5C, +siRNA TDP-43). After dox-induction, all three NLS mutant cell lines showed a similar cytoplasmic mislocalization of EGFP-TDP-43 in the absence of any stress (Fig. S5D). Upon treatment with sodium arsenite, a known inducer of oxidative stress, only the K82A and 83AAA mutants showed strong cytoplasmic EGFP-TDP-43 condensates (Fig. 5A). High resolution imaging after co-staining with an antibody for the SG marker G3BP1 revealed close proximity, yet no colocalization of EGFP-TDP-43 K82A and 83AAA with the G3BP1 signal, but a de-mixed phenotype, as recently described by Yan et al (Fig. 5A, see zoomed region)^29^. In contrast, while SGs were successfully formed in the EGFP-TDP-43 RK6A mutant, these cells only rarely formed cytoplasmic EGFP-TDP-43 condensates after 1h of arsenite treatment (Fig. 5A), corroborating our *in vitro* finding and demonstrating that this mutant is also strongly condensation-deficient in cells. Next, we tested whether the arsenite-induced cellular EGFP-TDP-43 condensation is linked to reduced solubility. To this end we performed RIPA extraction of either untreated or 1h arsenite-treated HeLa EGFP-TDP-43 NLS mutant cell lines and quantified the amount of RIPA-soluble and -insoluble TDP-43 by Western blot. In untreated cells, EGFP-TDP-43 WT similar to all NLS-mutants was completely soluble (top panel Fig. 5B). However, after arsenite treatment, solubility of TDP-43 WT, K82A and 83AAA was strongly reduced (Fig. 5B bottom panel, see C for quantification). In contrast, while solubility of EGFP-TDP43 RK6A was also reduced compared to the non-treated condition, it remained significantly more soluble than TDP-43 WT, K82A or 83AAA, further supporting an important role of the basic NLS residues in the condensation and solubility behavior of TDP-43.

**Figure 5:**
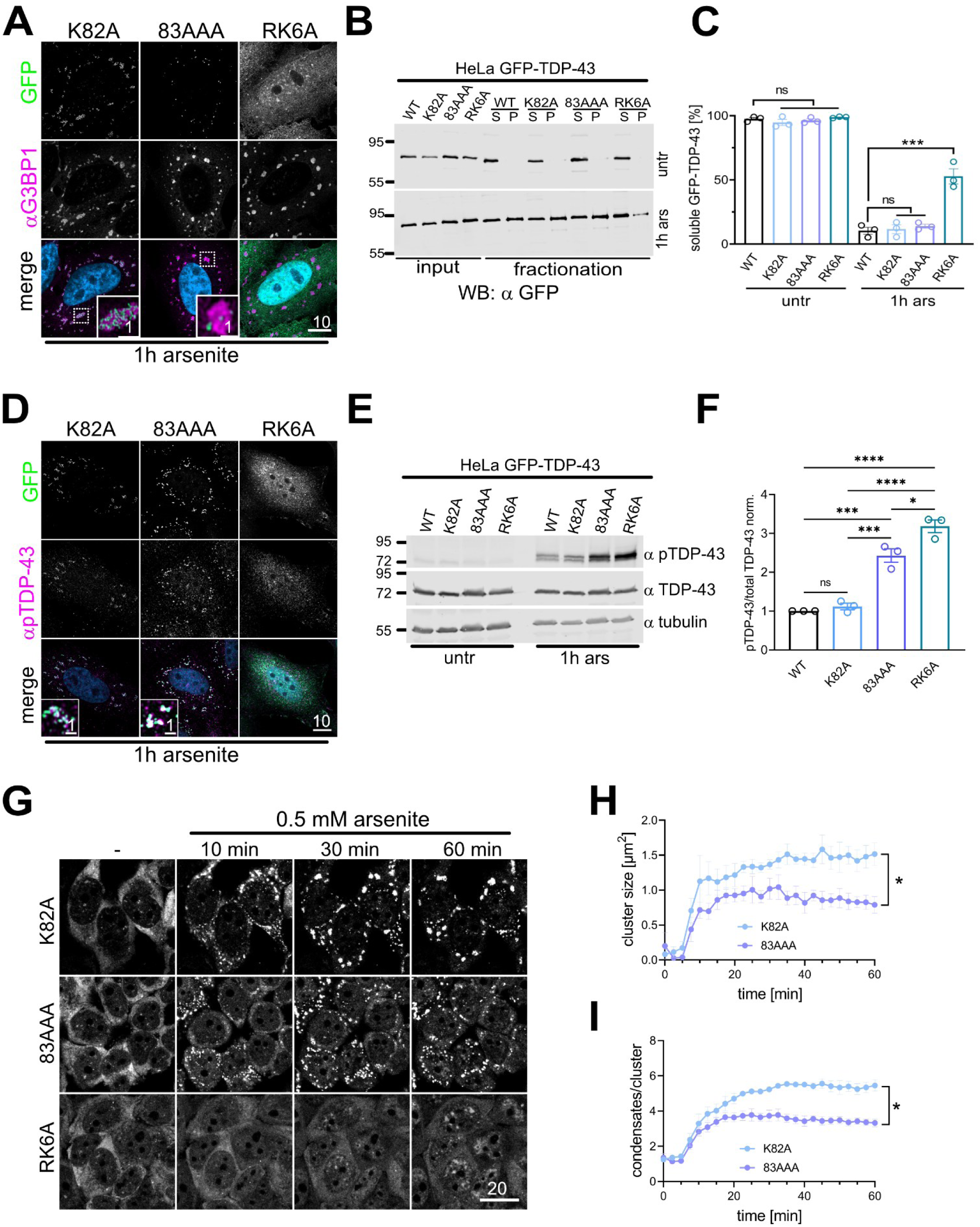
Cytosolic condensation and reduced solubility of TDP-43 upon arsenite stress depend on extent of basic-amino acid substitutions. A) Immunostaining for G3BP1 in HeLa cells stably expressing GFP-TDP-43 K82A, 83AAA or RK6A upon 1h treatment with 0.5 mM arsenite using Airyscan Confocal microscopy. Dashed boxes around SGs indicate zoomed-in region. Note, that for visibility the diffuse GFP-signal of TDP-43 RK6A was enhanced more strongly than for K82A and 83AAA, but identical to the GFP-signal in Figure S5. (B, C) Anti-GFP Western Blot (B) assessing RIPA-solubility of GFP-TDP-43 variants in absence or presence of arsenite stress. C) Quantification of percentage of soluble TDP-43 calculated as [S/(S+P)] for 3 independent replicates quantification, mean ± SEM, ns = non-significant, *** < 0.00021 by 1-way Anova with Dunett’s multiple comparison test to WT, untreated (untr) or 1h arsenite (ars) treated, respectively. D) phosphor-Ser409/410 TDP-43 (pTDP-43) immunostaining on HeLa GFP-TDP-43 K82A, 83AAA or RK6A cells upon 1h treatment with 0.5 mM arsenite. Dashed boxes indicate zoomed-in region. Note that for reasons of visibility, diffuse GFP signal for TDP-43 RK6A has been enhanced more strongly than condensed for K82A and 83AAAE (but equally to untreated cells). F) Western Blot analysis for levels of pTDP-43 (Ser409/410) versus total TDP-43 and α-tubulin of HeLa GFP-TDP-43 WT, K82A, 83AAA or RK6A cells either untreated (untr) or upon 1h treatment with 0.5 mM arsenite. F) Quantification of ratio of pTDP-43 to total TDP-43 normalized to WT of three independent experiments, mean ± SEM, ns = non-significant, * < 0.0332, *** < 0.00021, **** < 0.0001 by One-way Anova with Tukey’s multiple comparisons test. G) Representative still imagess from time-lapse imaging of HeLa GFP-TDP-43 stable cell lines carrying the indicated NLS mutations upon addition of 0.5 mM Na-arsenite. H, I) Analysis of average physical cluster size (H) and number of condensates per cluster (I) over time for 4 independent replicates with >68 cells each, mean ± SEM, * <0.021 by one-tailed Mann Whitney t-test for area under the curve.

C-terminal phosphorylation of TDP-43 is a hallmark of pathological TDP-43 aggregates in ALS and FTD patients^58,107–109^, which is recapitulated in the rNLS TDP-43 mouse model^95^ (corresponding to our 83AAA mutant), or upon arsenite stress in cell lines carrying the 83AAA NLS mutation^24,29^. To see if our K82A model also recapitulates this important feature of cytosolic TDP-43 pathology, we stained our EGFP-TDP-43 NLS mutant cell lines with a Ser409/410 phosphorylation-specific antibody (pTDP-43). Indeed, the arsenite-induced cytosolic condensates formed from EGFP-TDP-43 K82A and 83AAA colocalized strongly with pTDP-43 staining, while the RK6A line showed weak, diffuse phospho-S409/410 signal in the cytosol (Fig. 5D) compared to untreated cells (Fig. S5E). To obtain a more quantitative readout for TDP-43 phosphorylation in our EGFP-TDP-43 cell lines, we analyzed total cell lysates of untreated and arsenite-treated cells by Western blotting using the phospho-Ser409/410 antibody. In comparison to untreated cells, all EGFP-TDP-43 variants showed increased S409/410 phosphorylation of EGFP-TDP-43 upon 1h arsenite treatment (Fig. 5E). Surprisingly, the phospho-Ser409/410 levels for EGFP-TDP-43 83AAA and RK6A were significantly higher than for WT and K82A (Fig. 5E; see F for quantification), indicating that at least in our cellular model, TDP-43 K82A is the closest to WT in terms of self-assembly and phosphorylation behavior.

To compare the cytosolic condensation behavior of our HeLa EGFP-TDP43 NLS mutant cell lines to oxidative stress in more detail, we performed high resolution live cell imaging upon addition of arsenite and analyzed condensate formation over time. As seen before in fixed cells (Fig. 5A, D), EGFP-TDP-43 K82A and 83AAA strongly formed cytosolic condensates upon addition of arsenite, while RK6A stayed diffuse (Fig. 5G). Interestingly, for TDP-43 K82A, the cytosolic condensates appeared to cluster into significantly larger structures compared to TDP-43 83AAA, both in terms of physical cluster size (Fig. 5H) and number of individual condensates per cluster (Fig. 5I). Importantly, this distinct condensation behavior for K82A and 83AAA correlates well with the phase separation behavior we observed using recombinant proteins (Fig. 4), suggesting that it reflects the intrinsic condensation properties of the respective protein.

In summary, our data suggest that the minimalistic NLS mutant K82A represents the best model system for studying the cytosolic condensation and aggregation behavior of TDP-43 and thus is the preferred mutation of choice for cell and animal models used to recapitulate cytosolic TDP-43 pathology.

## Discussion

Aberrant phase separations have been linked to the pathological aggregation of TDP-43 in neurodegenerative disease^29,53,110–112^. To study TDP-43 aggregation, cellular and animal model systems often utilize TDP-43, in which the NLS has been mutated to render TDP-43 cytoplasmically mislocalized^24,29,92–98^. By performing phase separation experiments with recombinant proteins, we now find that either deleting the NLS-region (aa 79-101) or mutating the basic amino acids in TDP-43’s bipartite NLS involved in binding to Impα/β (K82/R83/K84/K95/K97/R98) strongly interferes with TDP-43 self-assembly. Importantly, this affects TDP-43 assemblies across size scales, i.e. mesoscale clusters at subcritical concentrations, µm-sized condensates visible by light microscopy, and detergent-resistant aggregates. This suggests that models using those mutants may not faithfully recapitulate TDP aggregation and thereby disease mechanisms.

In detail, we find that the disordered nature of the NLS-region contributes to, but is not sufficient to restore TDP-43 phase separation *in vitro*, as a replacement of the NLS-region with a disordered, non-charged linker still has a higher onset of condensate formation than TDP-43 WT. Surprisingly, deletion of the entire NLS-region (aa 79-101) or mutation of all five lysines in the NLS-linker region (K5A) impair TDP-43 dimer formation under non-phase separating conditions. So far, dimerization of TDP-43 is known to be suppressed by mutations or post-translational modifications (PTMs) of critical NTD residues, including amino acids located in or near the dimer interface, structurally relevant residues (L27/29) or deletion of the extreme N-terminal ten amino acids^6,36,98,113^. Mutations in the NLS-linker region, to our knowledge, were so far not known to influence TDP-43 dimerization. Notably, part of the NTD dimerization interface lies close to the NLS-region and includes K79^36^.

Even though mutating K79A alone does not render TDP-43 phase separation-deficient, additional mutations in the NLS-region (e.g., additionally mutating four lysines in the NLS or deleting the entire linker region) impair dimerization of full-length TDP-43. Our NMR data for the N-terminal fragment (aa 1-266) also indicate that the RK6A mutation structurally influences the NTD. This is in line with a recent study describing that, at least at acidic pH, the presence or absence of the NLS-region impacts the structure of the NTD^114^. However, in the context of the full-length protein and slightly basic pH, the RK6A mutation appears to have no significant effect NTD-mediated dimerization, as full length MBP-tagged TDP-43 RK6A behaves very similar to TDP-43 WT in size-exclusion chromatography, indicating a stable dimer formation of the RK6A mutant. Thus, even though the RK6A mutant in all phase separation assays behaved similarly to the 6M mutant, it is unlikely that its phase separation deficiency is caused simply by its inability to dimerize. Instead, our MD simulations indicate that basic residues in the NLS-region engage in diverse inter-chain interactions with various regions of TDP-43 within condensates. These interactions involve mainly aromatic residues in the C-terminal LCD, in particular tryptophan residues.

The C-terminal LCD alone also phase separates, and the α-helical CR as well as aromatic residues are known to be crucial for phase separation of the LCD^37,38,40,90^. Here, we demonstrate that condensation of full-length TDP-43 is very strongly decreased by mutation of all three tryptophan residues in the LCD (W3A: W305/385/412A), supporting our MD simulations. In contrast, mutation of the six phenylalanines (F6A: F267/283/289/367/397/401A), which were previously shown to strongly suppress phase separation of the LCD^90^, have only a minor effect on phase separation of full-length TDP-43. The strong condensation defect observed for the W3A mutant could in principle be explained by one of the mutated tryptophans being located in the α-helix of the CR region. Surprisingly, however, deletion of the entire CR only slightly delays onset of phase separation for full-length TDP-43, indicating that helix-helix interactions only play a minor role in the context of the full-length protein, in stark contrast to the LCD alone^37,38^. Instead, phenylalanine substitutions and deletion of the CR result in formation of larger TDP-43 condensates with pronounced wetting effect, which is in line with a reduced occurrence of π-π interactions.

In summary, our data clearly show that different and/or additional molecular interactions contribute to condensation of full length TDP-43 in comparison to the LCD alone, suggesting that TDP-43 condensates are multidimensional ensembles formed by a diverse set of inter-chain interactions of differential relevance: Consistent with a previous study^46^, we find that N-terminal dimerization is required for TDP-43 phase separation *in vitro*, yet is not sufficient. To form a network of contacts underlying condensate formation, the basic patches in the disordered NLS-region are essential, most likely by engaging in heterotypic interactions with aromatic (in particular Trp) residues in the LCD and possibly other regions, such as RRM1 and the NTD. Further LCD-LCD contacts, such as CR-CR or F-F interactions, likely modulate the nature of these condensates and only play a dominant role in the absence of the N-terminal regions. A role of diverse interdomain interactions for TDP-43 condensation has also been suggested by previous MD simulations^41,47^, even though interactions involving the disordered NLS-region were mainly found to be electrostatic (between R/K and D/E)^41^.

In contrast to micron-scale condensate formation, we see an equally strong impact of our various mutants on TDP-43 cluster formation at subcritical concentrations, indicating a different interaction hierarchy for cluster assembly. Our observations for cluster formation are in line with atomistic simulations by Pappu and colleagues who recently demonstrated that formation of microphases as precursors of clusters are driven by a contribution of homotypic (NTD-NTD or LCD-LCD) as well as heterotypic (LCD with all other domains) interactions^51^. While our clusters for TDP-43 at 0.7 µM detected by DLS (around 200 nm) exceed the size of microphases (30 nm), it is possible that such microphases are obscured by larger clusters in our DLS measurements, and/or that our recombinant TDP-43 has a different threshold concentration for microphase separation.

Our findings regarding the central role of the basic amino acids in the NLS for TDP-43 phase separation have important implications for model systems used to study cytosolic TDP-43 condensation and aggregation. Here we compared the phase separation propensities of two mutants widely used in cell and animal models, namely RK6A and 83AAA (K82/R83/K84A), with a TDP-43 variant in which only K82 is mutated (K82A) and which so far has not been used in disease models. We find that the K82A mutation is sufficient to inhibit nuclear import and cause cytosolic mislocalization of TDP-43 in cells, in line with previous reports^99,100^. Importantly, the three different NLS mutants exhibit a gradual deficiency to undergo cluster and condensate formation, correlating with the number of mutations. Only the K82A mutant forms condensates comparable to WT, while the 83AAA mutant shows already a stronger deficiency to self-assemble. Notably, also K82A showed slight defects in self-assembly at sub-critical concentrations, indicating that physiological processes that depend on TDP-43 cluster formation might already be compromised, yet this deficiency is clearly less pronounced than for the other two NLS mutants.

Our data also demonstrate that intrinsic phase separation properties of the three NLS mutants are largely conserved in the cellular context: The RK6A mutant is not only deficient in condensate formation *in vitro*, but also does not associate with SGs in semi-permeabilized cells and is strongly impaired in formation of cytoplasmic condensates upon short-term sodium arsenite treatment in intact cells. In contrast, both the K82A and 83AAA mutant still form cytoplasmic condensates upon arsenite treatment reminiscent of those recently described to arise upon oxidation-induced aggregation of TDP-43 by demixing from SGs^29^. While the RK6A mutant remains completely soluble and does not aggregate in our aggregation assay *in vitro* or upon arsenite treatment in cells, the K82A and 83AAA variants formed detergent-insoluble aggregates similarly to WT, both in vitro and in cells. Interestingly, however, the arsenite-induced cytoplasmic condensation behavior still differs between the K82A and 83AAA variants: TDP-43 K82A condensates associate into larger clusters consisting of more numerous smaller foci compared to TDP-43 83AAA, indicating that TDP-43 K82A reacts more sensitively to cellular stress than the triple alanine mutant.

Phosphorylation is a pathological hallmark of cytosolic TDP-43 aggregates^58^, and arsenite-induced cytoplasmic condensates for both EGFP-TDP-43 K82A and 83AAA are stained with an antibody specific for Ser409/410 phosphorylation, suggesting that they can form pathology-related assemblies in cells. Unexpectedly, quantification of TDP-43 phosphorylation levels by immunoblot revealed a significantly higher level of phospho-Ser409/410 for TDP-43 RK6A and 83AAA compared to WT and K82A upon arsenite treatment. A possible explanation for this observation could be that more extensive mutations in the NLS render TDP-43 into a primed substrate, resulting in higher phosphorylation levels of the S409/410 site or even across the entire length of TDP-43. Alternatively, the LCD in the 83AAA and RK6A variants could be more accessible, for example to kinases, due to the reduced condensation properties of these variants. It remains to be seen if additional phosphorylation sites, or even other PTMs, show increased levels in the more extensively mutated 83AAA and RK6A variants and how this compares to the TDP-43 PTM pattern in ALS/FTD patients. It will also be interesting to investigate whether more or different cellular proteins co-aggregate with TDP-43 K82A compared to the 83AAA variant.

In conclusion, our work presented here strongly suggests that K82A and 83AAA NLS mutants are better suited cellular model systems than the RK6A mutant for studying cytosolic TDP-43 phase separation and aggregation. TDP-43 K82A most closely reflects the self-assembly behavior of TDP-43 WT, and pathological hallmarks, such as S409/410 phosphorylation levels, are most closely mimicked. Hence, we suggest that TDP-43 K82A is the preferred NLS mutant of choice in cellular or animal models to study TDP-43 aggregation in the cytoplasm.

## Acknowledgements

The authors thank all members form the Dormann group for critical discussion. The authors gratefully acknowledge S. Möckel and S. Nick at IMB flow-cytometry core facility for assistance with operating the Bigfoot cell sorter instrument (project number 511658729). We acknowledge support by the CRC1551 internal service projects Z01 (Biopolymer Engineering and Bioanalytics at IMB’s Protein Production Core Facility) and Z02 (Advanced Imaging at JGU Bio microscopy core facility and IMB’s Microscopy Facility) and we thank the facility staff of the IMB Protein Production and the IMB Microscopy Core facilities for technical support. We further thank Elmar Jaenicke from the JGU Biomolecules and Bioanalytics Core Facility for his support with DLS measurements. We thank Marc-David Ruepp (Kings College, London, UK) and Christian Behrends (LMU Munich, Germany) for HeLa and Hela Flp-In cells, respectively. T.M. thanks the Center for Medical Research, Medical University of Graz, Graz, Austria for laboratory access, as well as Pedro Musso and Johanna Pirchner for their assistance in protein expression and purification.

This project was funded by the Deutsche Forschungsgemeinschaft (DFG, German Research Foundation) - CRC1551 – Project No. 464588647. D.D. furthermore acknowledges funding from the European Union (EU) through the ERC CoG TDP-ASSEMBLY (Project 1011258709), the DFG within the Heisenberg Programme (project ID 442698351) and the SPP2191 “Molecular Mechanisms of Functional Phase Separation” (project ID 419139133). X.P. acknowledges a PhD fellowship from the Max Planck Graduate Center (MPGC) Mainz. X.P. was funded by the DFG, Project No. 233630050 – TRR 146. L.S.S. and X.P. thank M3ODEL for support. L.S.S and D.D. gratefully acknowledges support from ReALity (Resilience,Adaptation and Longevity) and Forschungsinitiative des Landes Rheinland-Pfalz. T.M. is grateful to the Austrian Science Fund (FWF) for excellence cluster 10.55776/COE14, Grants DOI 10.55776/P28854, 10.55776/I3792, 10.55776/DOC130, and 10.55776/W1226, the Austrian Research Promotion Agency (FFG) grants 864690 and 870454; the Integrative Metabolism Research Center Graz; the Austrian Infrastructure Program 2016/2017; the Styrian Government (Zukunftsfonds, doc.fund program); the City of Graz; and BioTechMed-Graz (flagship project). This project was funded in part by the FFG and the European Union (EFRE) under grant 912192. The Spinning Disk Confocal System (VisiScope 5-Elements, IMB Microscopy Core Facility) was supported by the DFG (DFG Project Number 402386039). The ZEISS LSM 900 was funded by the Ministry of Science and Health of Rhineland-Palatinate and the European Regional Development Fund (ERDF/REACT-EU, Grant No. 84012490).

---

**Figure S1 (to Figure 1):**
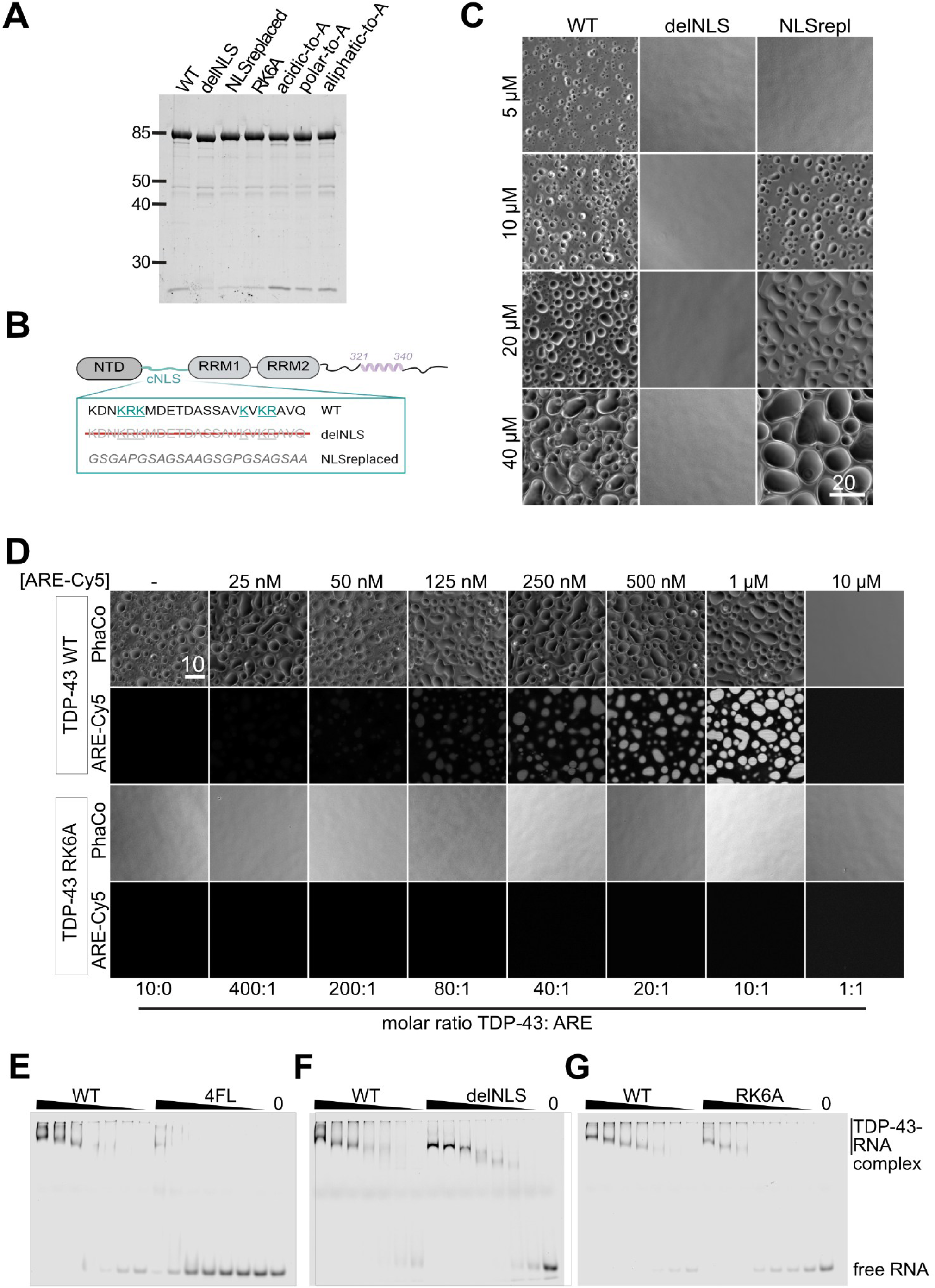
The disordered NLS-region of TDP-43 is required for its phase separation, but not for its RNA-binding. A) Sypro-Ruby stained SDS-PAGE showing similar purity for recombinant TDP-43-tev-MBP-His_6_ versions (∼ 445 ng/lane). B) Cartoon of TDP-43 WT and TDP-43 mutant with the NLS-region (aa 79-101) either deleted (delNLS) or replaced by a disordered, non -charged linker of identical length (NLSrepl). C) Representative phase contrast images for TDP-43 WT, delNLS and NLSrepl condensates at different concentrations. Bar, 20 µm. D) Representative images of condensates formed by 10 µM TDP-43 WT or RK6A, respectively, after 1h Tev protease cleavage in presence of increasing amounts of Cy5-labelled TDP-43 autoregulatory element (ARE) RNA. Cy5-Fluorescence images were acquired using structural illumination. E-G) EMSAs for TDP-43-tev-MBP-His_6_ with IR700-labelled ARE-RNA demonstrating strongly reduced RNA-binding capacity of RNA-binding deficient TDP-43 4FL (F147/149/229/231L) mutant (E), while both TDP-43 delNLS (F) or TDP-43 RK6A (G) show similar RNA-binding capacity as WT.

**Figure S2 (to Figure 2):**
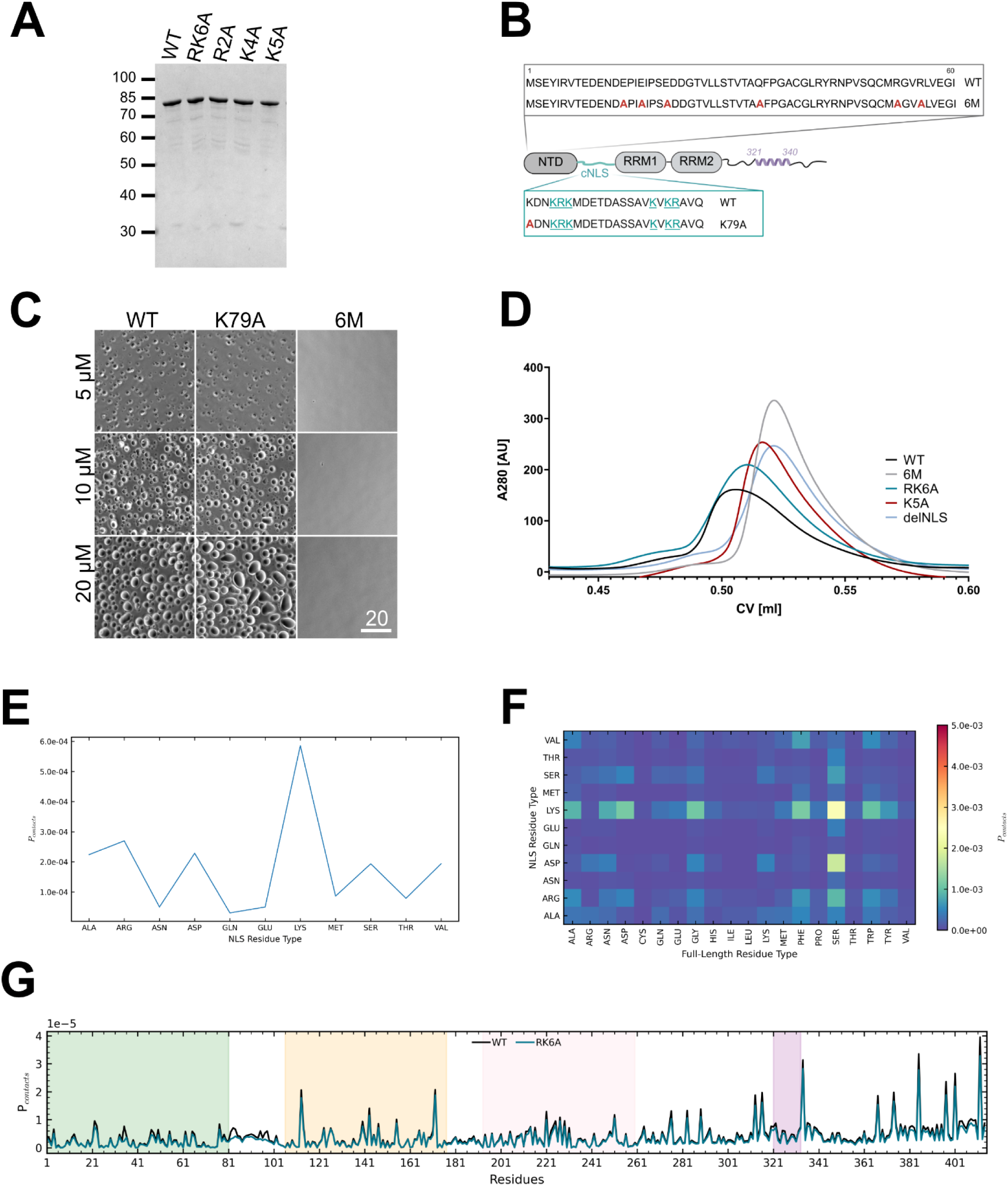
High number of lysines in the disordered NLS-region is required for TDP-43 phases separation. A) Sypro-Ruby stained SDS-PAGE showing similar purity for recombinant TDP-43-tev-MBP-His_6_ versions used in Figure 2 (∼ 445 ng/lane). B, C) Cartoon (B) showing mutations for multimerization deficient 6M and K79A mutant of TDP-43 used in (C) to analyze their phase separation propensity by phase contrast microscopy. (D) The indicated purified TDP-43-MBP-His variants were subjected to size-exclusion chromatography analysis monitored at 280 nm (A280) to address their dimerization capability in comparison to dimerization–competent (WT) or deficient (6M) TDP-43. E) Non-normalized contact frequency of residue types within the disordered NLS-region against residues in full-length protein chains F) Non-normalized contact probability of residues disordered NLS-region over full-length TDP-43. G) Comparison of the residue inter-chain contact possibility of TDP-43 WT (n=3 independent replicates of ∼28.3 ± 2.3 µs simulation time) and RK6A (single replicate of 37 µs). Folded regions are indicated in green (NTD), yellow (RRM1), rose (RRM2) and purple (helix in LCD).

**Figure S3 (to Figure 3):**
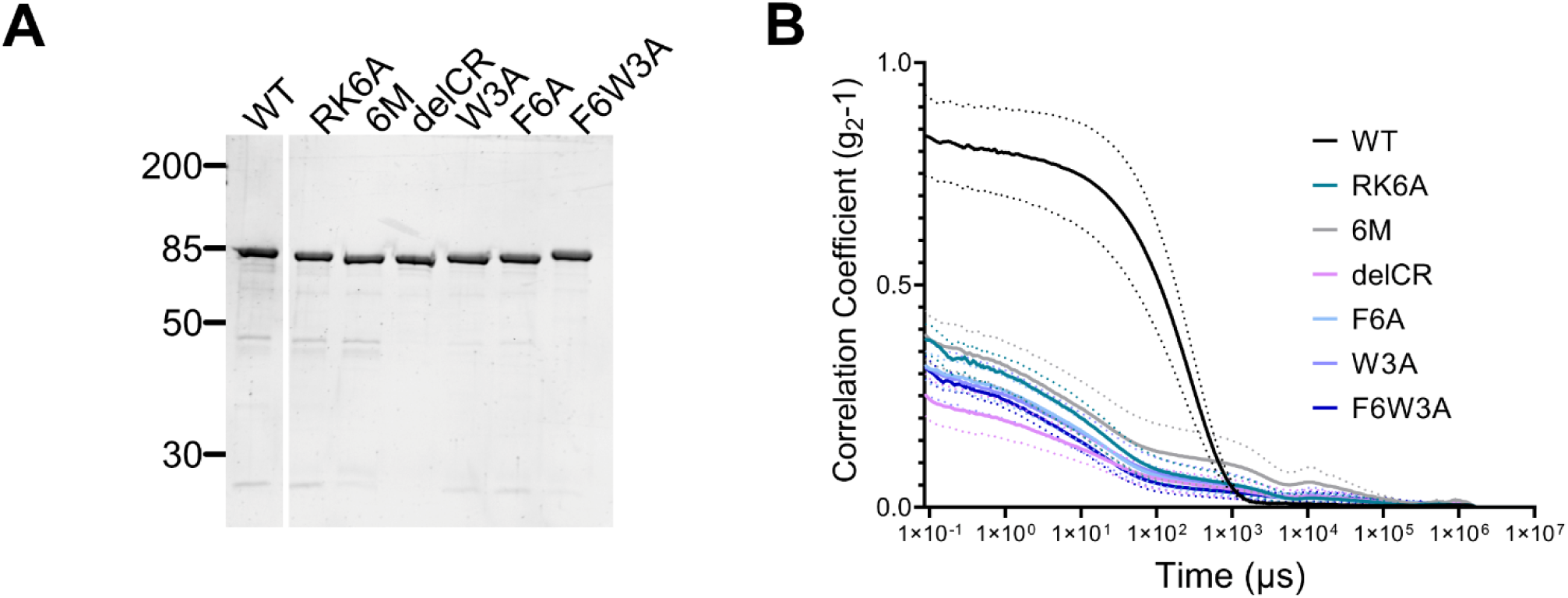
Interactions of the NLS-region with C-terminal aromatic residues are important for both condensation and nanocluster formation of TDP-43. A) Sypro-Ruby stained SDS-PAGE showing similar purity for recombinant TDP-43-MBP-His versions used in Figure 3 (∼ 445 ng/lane). Note, one lane was spliced out to remove one mutant not shown in Figure 3. B) Correlation curves of DLS measurements underlying intensity profiles shown in Figure 3 E, F. Note, all mutants show poor scattering compared to WT caused by reduced nanocluster formation. Mean of 3 independent experiments ± SEM indicated by the dotted lines.

**Figure S4 (to Figure 4):**
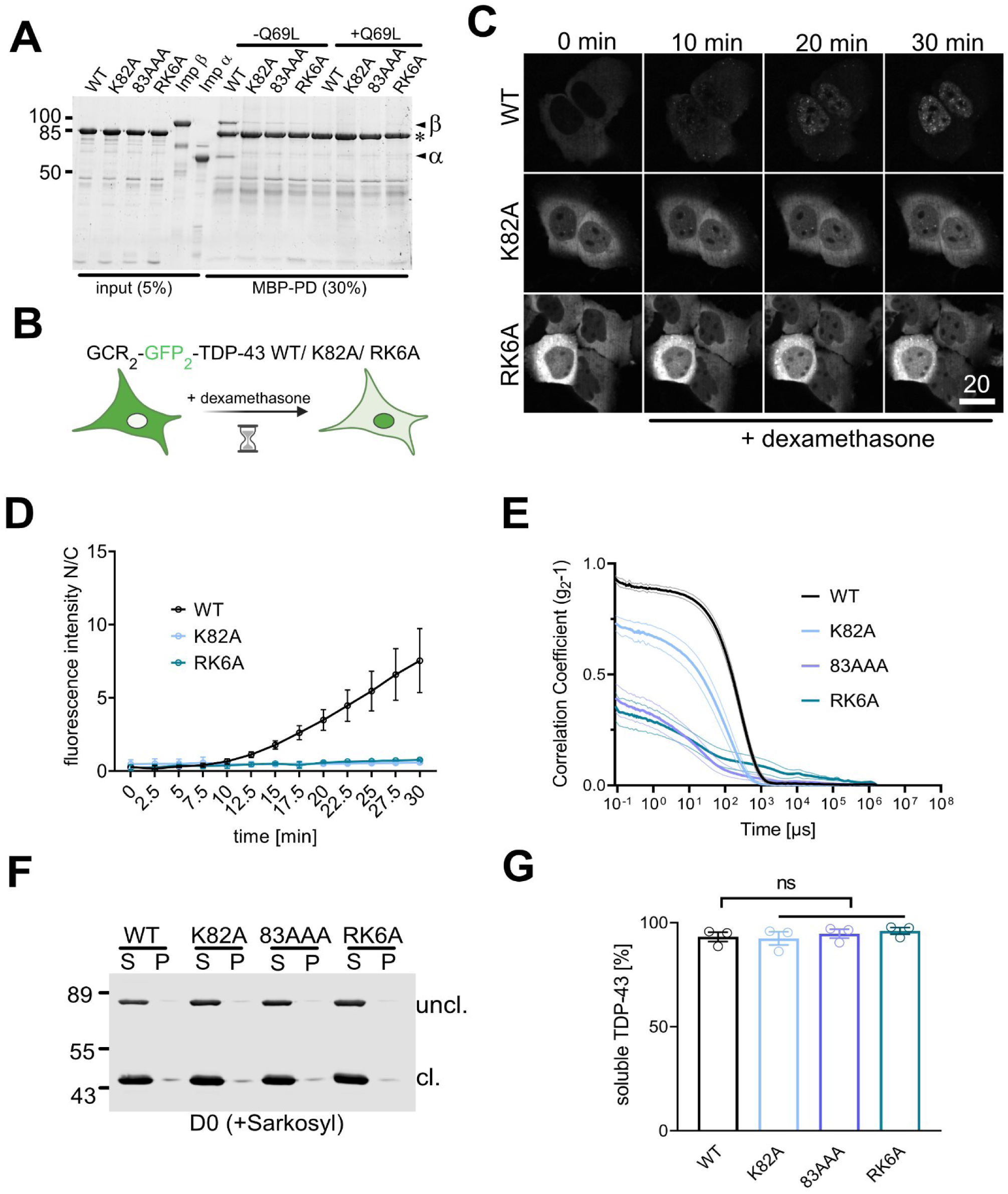
Differential phase separation and aggregation behavior of commonly used TDP-43 import-deficient mutants in vitro. A) MBP-pulldown for TDP-43 WT and indicated NLS-mutants with Impα3/β in presence or absence of RanQ69L-GTP. The bait (TDP-43-tev-MBP-His_6_) is marked by an asterisk, Impα (α) and Impβ (β) indicated by arrowheads. B) Scheme of hormone-inducible import assay using GCR_2_-GFP_2_-TDP-43. C) Representative images (C) and quantification (D) from time-lapse recordings of HeLa cells transiently transfected with indicated GCR_2_-GFP_2_-TDP-43 reporters upon addition of doxycycline. D) Quantification of relative nuclear import quantified as fluorescence intensity in the nucleus (N) over cytoplasm (C) over time. Mean ± SD of 2 (K82A) - 3 (WT, RK6A) independent experiments of at least 40 cells each. E) Correlation curves of DLS measurements underlying intensity profiles shown in Figure 4E. Note, reduced scattering by K82A as well as 83AAA or RK6A mutant, respectively, compared to WT caused by reduced self-assembly of those mutants. Mean of 3 independent experiments ± SEM indicated by the dotted lines. F, G) Sedimentation analysis after sarkosyl extraction in the aggregation assay at day 0 (D0) analyzed by Western Blot (F) using an N-terminal antibody. G) Quantification of 3 independent replicates of the sedimentation analysis coupled to sarkosyl extraction at D0 (mean ± SEM). ns, non-significant.

**Figure S5 (to Figure 5):**
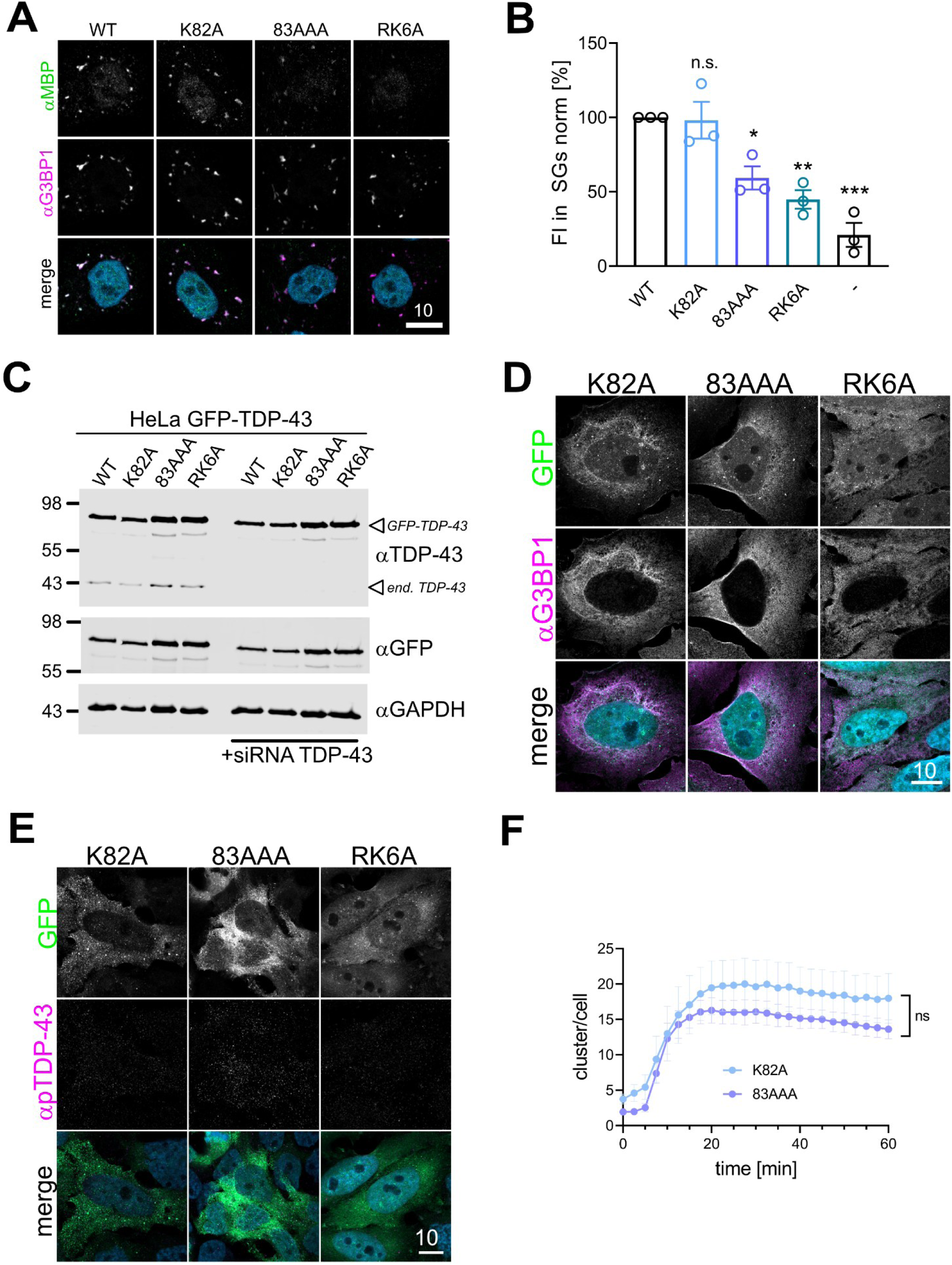
Cytosolic condensation and reduced solubility of TDP-43 upon arsenite stress depend on extent of basic-amino acid substitutions. (A) Representative confocal fluorescence microscopy images of a SG association assay in semi-permeabilized HeLa cells for the indicated TDP-43-tev-MBP-His_6_ NLS variants. (B) Quantification of the respective association of the individual TDP-43 variants normalized to WT as mean of three independent experiments ± SEM; ns, not significant, *<0.021, **< 0.0021, ***<0.0002by 1-way ANOVA with Dunett’s multiple comparison test compared to WT. (C) Western Blot of cell lysates of the indicated stable HeLa GFP-TDP-43 FlpIn lines in absence or presence of endogenous TDP-43 KD (+siRNA TDP-43) with antibodies against TDP-43, GFP or GAPDH (as loading control). (D) G3BP1 immunostaining of unstressed HelaFlpIn GFP-TDP-43 K82A, 83AAA and RK6A, respectively. Note, that for visibility, the signal for GFP and G3BP1 (AF647) was enhanced more strongly than in the main figure. (E) phospho - Ser409/410 TDP-43 (pTDP-43) immunostaining on untreated HeLa GFP-TDP-43 K82A, 83AAA or RK6A cells. Please note that for reasons of visibility, the diffuse GFP signal was enhanced more strongly than in the main figure for K82A and 83AAA upon arsenite (but equally to RK6A + ars). (F) Analysis of average number of clusters in HeLa GFP-TDP-43 K82A and 83AAA cells, respectively, upon Na-arsenite treatment over time for 4 independent replicates with >68 cells each, mean ± SEM, * <0.021 by two-tailed Mann Whitney t-test for area under the curve.

## Notes

### Competing Interest Statement

The authors have declared no competing interest.

